# Direct Assessment of Short-Latency Intracortical Inhibition via Immediate TMS-Evoked Potentials

**DOI:** 10.64898/2026.04.15.718740

**Authors:** Lasse Christiansen, Yufei Song, Ditte Haagerup, Mikkel Malling Beck, Kora T. Montemagno, John Rothwell, Hartwig Roman Siebner

## Abstract

Short-interval intracortical inhibition (SICI) is the most widely used neurophysiological index of GABAergic inhibition in the human cortex. However, it is an indirect measure, inferring synaptic inhibition from suppression of peripherally recorded motor-evoked potentials (MEPs) elicited by transcranial magnetic stimulation (TMS). In the standard protocol, a subthreshold conditioning pulse suppresses the MEP evoked by a suprathreshold test pulse delivered 1–5 ms later. Interpretation is further complicated by temporal overlap with short-interval intracortical facilitation (SICF), reflecting excitatory interactions at interstimulus intervals of ∼1.5 and 2.7 ms.

To overcome these limitations, we recorded immediate TMS-evoked EEG potentials (iTEPs; 1–10 ms post-stimulus) as a more direct measure of motor cortical activity in 16 healthy volunteers (20–35 years; 7 male). The conventional SICI protocol suppressed only later components of the iTEP, likely corresponding to late corticospinal volleys previously identified in epidural spinal recordings after suprathreshold TMS, while the earliest iTEP component was unaffected. Importantly, later iTEPs were suppressed to a similar extent whether conditioning–test intervals coincided with SICF peaks or troughs, and the magnitude of iTEP suppression correlated with concurrently recorded paired-pulse MEP suppression. SICI also reduced an early TEP component (N15; 10–20 ms), but paired-pulse N15 suppression showed a different dependence on stimulus intensity and did not correlate with MEP suppression.

These findings demonstrate that SICI measured via MEPs does not reflect a global index of cortical GABAergic motor cortical inhibition but instead reflects inhibition within specific cortical circuits that can be investigated directly with iTEPs.

## Introduction

Local cortical inhibition is thought to be essential for shaping the timing and spatial organization of descending corticomotor control [1,2]. In human motor cortex (M1), it is conventionally probed using paired-pulse inhibition of the motor evoked potential (MEP) using a technique known as short-latency intracortical inhibition (SICI). This is assessed in the hand representation of M1 (M1-HAND) where MEPs in contralateral muscles are suppressed when preceded by a subthreshold conditioning TMS pulse delivered 1–5 ms earlier [3]. Indirect evidence suggests inhibition is mediated by intracortical interneurons via GABA-A receptors [4,5]. As a result, SICI is often employed as a general measure of cortical GABAergic tone. For example, SICI is modulated by motor state [6], and reduced during motor learning [7], consistent with the disinhibition that enables learning-related plasticity in rodent motor cortex [8,9]. Clinically, reduced SICI has been used as a window into deficient inhibition focal dystonias [10] and amyotrophic lateral sclerosis [11]. However, SICI is a very indirect measure of inhibition in the cortex since it relies on the MEP, which is several synapses removed from the presumed inhibitory synapses in cortex. SICI is also difficult to assess in patients with damage in the corticomotor pathway. A method that can index SICI directly at the cortical level would therefore improve mechanistic interpretation and increase clinical utility.

Previous work using invasive recordings has shown that SICI is a more nuanced measure than appears from the study of MEPs. A single suprathreshold TMS pulse over the precentral gyrus evokes descending volleys in fast-conducting corticospinal motoneurons[12]. These volleys can be recorded invasively with epidural electrodes at cervical [13] and thoracic levels[14]. At peri-threshold intensities, the corticospinal volleys consist of one or more indirect (I)-waves separated by 1-1.5 ms, reflecting transsynaptic activation of corticospinal neurons. The number and size of the I-waves depend on stimulation intensity and the orientation of the induced tissue current relative to the precentral sulcus[15]. Although the precise circuitry generating I-waves remains debated [16], recent computational and experimental evidence suggests that they arise from population responses shaped by intracortical excitation and GABAergic inhibition [17]. Consistent with this view, SICI suppresses the amplitude and number of later I-waves in epidural recordings [18] but has little or no effect on the earliest volley.

A complication in interpreting SICI is that in addition to producing inhibition, paired-pulse TMS can also elicit short-latency intracortical facilitation (SICF) using the same range of interstimulus intervals (ISIs) employed in the paired-pulse SICI paradigm. SICF emerges when a slightly suprathreshold pulse precedes a slightly subthreshold pulse or when two slightly subthreshold pulses are paired, leading to MEP facilitation at ISIs corresponding to I-wave periodicity [17–19]. With biphasic pulses, the first two SICF peaks typically occur at ISIs of ∼1.5 and ∼2.7, with a trough at ∼2.0. [20]. Because SICI and SICF occur within the same early temporal window, they may involve partially overlapping neural circuits. However, the evidence is mixed: some studies report complex interactions that depend on the amplitude of the response to the unconditioned test MEP [21–23], whereas others find comparable SICI at both, SICF peaks and troughs [24,25]. A major obstacle to resolving these discrepancies is the inability of conventional TMS to directly assess paired-pulse effects at the cortical level.

Epidural recordings are invasive and rare, but recent developments have made it possible for TMS-EEG to measure cortical responses directly in the human brain, and several studies have examined paired-pulse effects on TMS-evoked potentials (TEPs): for instance, Cash et al., (2016) reported bidirectional changes in the amplitude of the positive TEP peaks around 30 and 60ms following paired-pulse stimulation. However, these relatively late latencies (≥ 30 ms) leave ample time for contributions from subcortical, peripheral, and re-afferent inputs. As a result, conventional TMS-EEG offers only an indirect and mechanistically ambiguous view of early intracortical inhibition. Importantly, many accounts of mid-latency TEP suppression view it in the same way as MEP suppression; that is, as a global indicator of non-specific GABA inhibition at the stimulated site with the added advantage that it can be used in any area of cortex outside motor regions. However, this ignores the work described above on I-waves that suggests that SICI affects certain, but not all, cortical circuits, at least in patients with epidural electrodes implanted for intractable dorso-lumbar pain. Currently, no evidence for circuit-specificity of SICI exists in able-bodied humans.

Recently, our laboratory identified a fast EEG signature that aligns temporally with early TMS-evoked corticospinal activity [28]. Single-pulse stimulation of the M1-HAND region elicited immediate TMS-evoked potentials (iTEPs), a sequence of rapid peaks superimposed on a slower wave appearing∼2-6ms after stimulation [26,27]. The multi-peak iTEP mirrors the physiological properties of corticospinal I-waves: its peak number and relative peak amplitudes depend systematically on the intensity and orientation of the TMS-induced electrical field in M1-HAND.

Here, we tested whether iTEPs provide a direct non-invasive measure of intracortical inhibition in the human cortex. First, we assessed the effect of the paired-pulse SICI paradigm on iTEP amplitude and related paired-pulse effects on iTEP peak amplitudes to the conventional MEP-based SICI measure. We then examined whether paired-pulse iTEPs are modulated by SICF related mechanisms. Finally, we compared immediate and later EEG markers of inhibition.

## Methods

### Participants

Sixteen able-bodied participants (7 males) aged 20-35 years were included in the experiment (see Suppl. Table 2). All participants were screened for TMS contraindications, and all procedures were carried out in accordance with the Helsinki declaration and were approved by the regional ethical committee (Capital Region of Denmark; Protocol number H-15008824).

### Experimental procedures

Figure 1 depicts the experimental flow. During the main experimental day, we first identified the TEP optimized hotspot for the First Dorsal Interosseous (FDI) muscle where single-pulse TMS could be delivered with 110% resting motor threshold (RMT) without muscular artefacts following the same iterative procedure as previously reported [26]. After establishing RMT and Active Motor Threshold (AMT) at the TEP optimized hotspot, an individual SICF curve was established (Figure 1A). The interval corresponding to the trough between the first two SICF peaks was used for establishing the relationship between CS intensity and MEP inhibition for each participant (Figure 1B). The CS intensity that resulted in the most pronounced MEP suppression was used for the following paired pulse TMS-EEG-EMG recordings where ISIs were adjusted to the peaks and rough of the SICF-curve (Figure 1C). In four participants, a follow-up experiment investigated the effect of CS intensity on iTEP and TEP suppression (Figure 1D).

**Figure 1.**
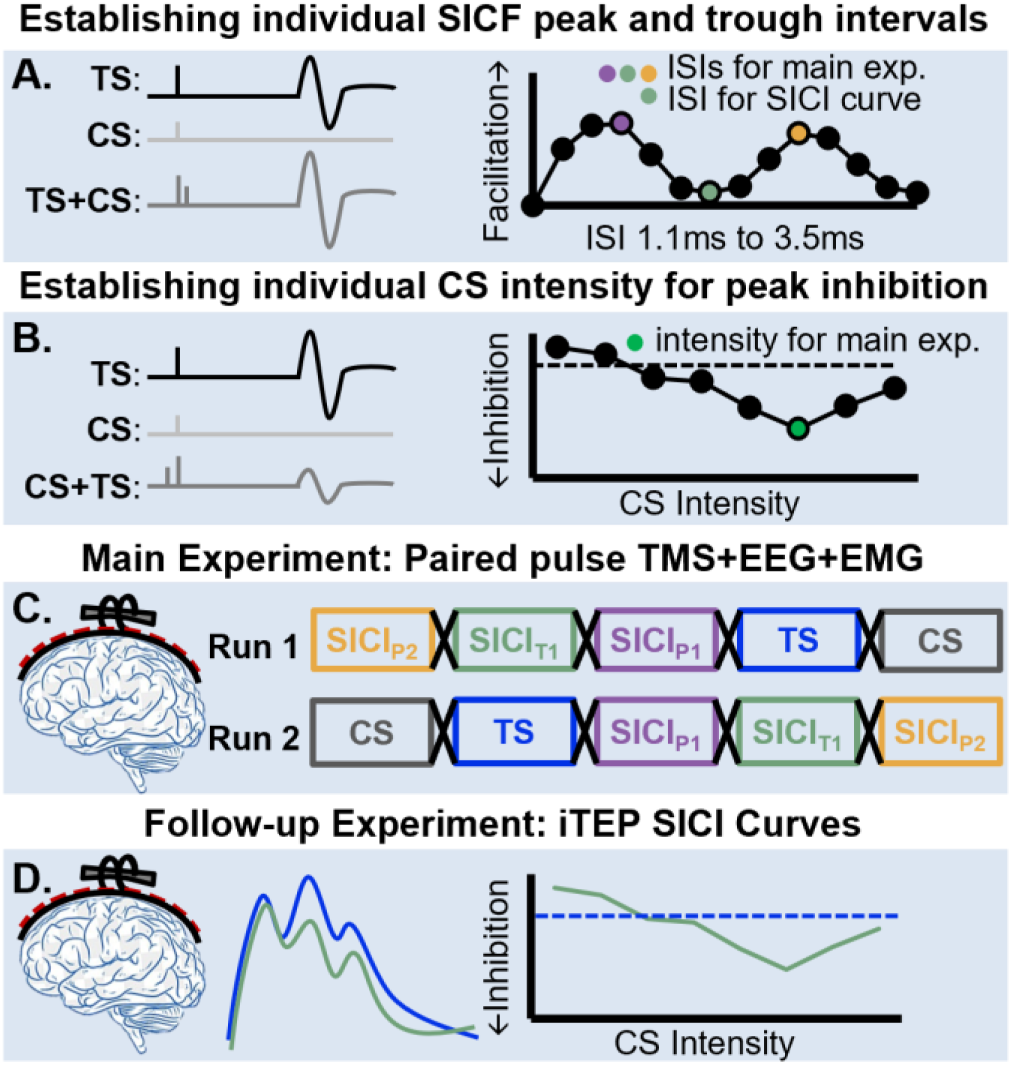
Experimental procedures. **A.** Individual Short-latency Intracortical facilitation (SICF) MEP profile was established, and the first peak, the first trough and the second peak were computed and extracted for later use. **B.** The interstimulus interval corresponding to trough 1of the SICF profile was used as ISI in a SICI recruitment curve aimed to delineate the CS intensity that resulted in the largest magnitude of inhibition of MEPs. **C.** Main experiment with TMS-EEG and EMG recording. Included 5 different conditions, 2 single-pulse and 3 paired –pulse conditions. Trials were recorded over two runs consisting of 50 trials of each condition. **D.** Follow-up experiment. In a small cohort, *n*= 4, 90 trials of single-and paired-pulse conditions were recorded with CS intensities between 40% and 90% AMT, to find the minimal CS intensity needed to suppress iTEP amplitude. AMT = Active Motor Threshold, CS = Conditioning Stimulus, P1= Peak 1, P2 = Peak2, SICI= Short-latency Intracortical Inhibition, T1= Trough 1, TS= Test Stimulus.

The CS intensity and interstimulus interval used for TMS-EEG-EMG recording were kept constant within-subject and established as described below.

Before recording, the participants were equipped with noise-masking consisting of pink noise with recorded TMS clicks generated using TAAC software titrated to fully mask TMS clicks delivered 5-10 cm above the scalp or to the highest tolerable intensity[28]. The TMS-EEG recoding was done in two runs each consisting of five recordings corresponding to five conditions; single-pulse test stimulus (TS), single-pulse conditioning stimulus (CS), paired-pulse stimulation with ISI adjusted to the first peak (SICI_P1_), the first trough (SICI_T1_) and the second peak (SICI_P2_) of the SICF curve. The five conditions were randomly ordered within run 1 and then reversed for run 2 to mitigate potential order effects within-subject. The order of conditions was pseudo-randomized and counterbalanced between-subjects.

### Transcranial Magnetic Stimulation

During all experiments, the TMS coil positioning was monitored via stereotaxic neuronavigation (TMS Navigator, Localite, GmbH, Bonn, Germany) using individual T1-weighted brain images obtained with a 3T MR scanner (PRISMA, Siemens, Erlangen., Germany or Achieva, Philips, Best, The Netherlands). Biphasic TMS pulses were delivered to the left M1-HAND using a 35 mm figure-of-eight coil (MC-COOL-B35-HO coil, MagVenture X100 with MagOption, MagVenture A/S, Farum, Denmark) with active cooling while the right hand was at rest. The recharge delay of the stimulator was set to 500 ms after the stimulation to avoid potential recharge artifacts in our EEG recordings. Furthermore, a thin layer of foam (approx. 1.5 mm thick) was placed between the coil and the electrodes to reduce bone conduction of sound and vibration-induced somatosensory co-activation. The active motor threshold (AMT) was defined as the minimum intensity needed to evoke a clear MEP in the tonically contracting right FDI muscle with a peak-to-peak amplitude above 200 µV followed by a silent period. RMT was defined as the minimum intensity to evoke an MEP with an amplitude above 50 µV, both in at least 5 of 10 trials during complete muscle relaxation [29].

### EEG and EMG recordings

EEG signals were recorded in BrainVision Recorder software (Brain Products GmbH, Germany) from 61 passive Ag/AgCl electrodes placed in an equidistant EEG cap (M10 cap layout, BrainCap TMS, Brain Products GmbH, Germany), with the reference and ground electrodes placed on the right and left side of the participants’ forehead, respectively. A TMS-compatible EEG amplifier (actiCHamp Plus 64 System, Brain Products GmbH, Germany) was used at a sampling frequency of 50 kHz (*anti*-aliasing low-pass filter at 10,300 Hz). All electrodes were prepared using electroconductive abrasive gel to reduce impedance (<5 kΩ), which was regularly checked during the experiment. During TMS-EEG recordings, participants were asked to fix their gaze at fixation spot approx. 80 cm in front of them, to fully relax, keep their eyes open, and avoid eye blinks in response to TMS.

Electromyographic (EMG) recordings were acquired from the right first dorsal interosseus (FDI, *n*=16) and Abductor Digiti Minimi (ADM, *n*=15 due to incomplete recording for one participant) in a belly-tendon montage with the ground positioned on the right ulnar styloid process. EMG signals were amplified (x500), bandpass filtered (10–2000 Hz), and sampled at 5000 Hz (Digitimer D360, Digitimer Ltd., Hertfordshire, UK) using Signal software (v4.17, Cambridge Electronic Design, Cambridge, UK).

### MEP measurements of Short-latency Intracortical Facilitation (SICF)

In all participants, SICF curves were obtained by delivering single and paired pulses with the subthreshold conditioning pulse (CS) after the suprathreshold test pulse (TS) every 5 s (20% jitter). We included one single-pulse (TS) and thirteen paired-pulse TMS conditions using ISIs ranging from 1.1 to 3.5 ms at a temporal resolution of 0.2 ms. Fifteen MEPs were recorded per stimulation condition, with conditions being pseudo - randomized within three blocks of 70 trials for a total of 210 trials. The TS was adjusted to 110 % RMT and the CS to 90% RMT. Individual SICF peaks and trough were determined. The intervals corresponding to the two first SICF peaks (P1 and P2) as well as the first trough (T1) were used to individually adjust ISIs during TMS-EEG-EMG. To accommodate the non-gaussian positively skewed distribution of MEP amplitudes with 110% RMT [30,31], MEPs were log-transformed, and the peaks and trough were identified from the highest and lowest average of log-transformed MEPs.

### MEP measurements of Short-latency Intracortical Inhibition (SICI) recruitment curves

The ISI corresponding to the T1 between P1 and P2 of the SICF curve was used to generate individual SICI recruitment curves. The effect of CS intensity on SICI was measured in steps of 5% AMT spanning an intensity range from 45% to 115% of AMT. In correspondence with previous research [32], five trials from each paired pulse conditions (given every 5s with 20% jitter) were collected successively with the order of conditions randomized. Five single-pulses were delivered before and after the paired pulse conditions (total of 10 single-pulses). The CS intensity that resulted in the largest MEP suppression was used for subsequent TMS-EEG recordings. Single pulse and TS intensity was 110% of RMT.

### TMS-EEG-EMG recordings – Main Experiment

One hundred trials were recorded for each of the five conditions (TS, CS, SICI_P1_, SICI_T1_, SICI_P2_) divided into two runs. Within each run blocks of fifty trials of each of the five conditions were recorded with an ISI of 2s and a jitter of 20%. Order of blocks were pseudo-randomized and counterbalanced between participants. To minimize potential effects of order or slow drifts within-subject, the order of conditions within run 2 was reversed from run 1. A small break without noise masking was provided between each condition along with a longer one between the two runs. Single pulse and TS intensity was 110% of RMT, whereas CS intensity was individualized as described above. In a few participants TS intensity was increased with a few percentage points to accommodate absent MEP responses during the recording of SICF- or SICI recruitment curve (see Table S2).

### TMS-EEG-EMG recordings – Follow-up Experiment

In four subjects, ninety trials were collected for each of the 6 paired pulse (SICI_40_, SICI_50,_ SICI_60,_ SICI_70,_ SICI_80,_ SICI_90_) and 5 single pulse (TS, CS_40_, CS_60,_ CS_80,_ CS_90_) conditions in 3 runs of 30 trials. Hotspots and thresholds were established as described for the main experiment, and the ISI was set to 2.1 ms (n=2) or 2.3 ms (n=2). Noise-masking was adjusted as described above.

### Data analysis

EEG and EMG data were preprocessed and analyzed in MATLAB (R2024b) using customized scripts and functions from EEGLAB [33] and Fieldtrip [34].

EMG trials were flagged for rejection when visual inspection identified background activity in the pre-stimulus interval (−100 to 0 ms). EEG data were epoched from −1000 to 1500 ms relative to the TMS pulse (for TS and CS) or the second pulse (for SICI_P1_, SICI_T1_, SICI_P2_) and baseline-corrected using −210 to −10 ms. To remove slow drift, we applied robust detrending and re-epoched the data into a shorter interval (-500 to 500 ms) to minimize edge artifacts [35] and remove any overlapping signal segments. EEG was then inspected trial-by-trial. Channels with excessive noise or long-lasting decay artifacts were excluded. Trials with prominent noise, large blinks, or eye movements were flagged for rejection. For each participant and condition, EMG- and EEG-based trial flags were combined (union) into a single rejection mask, which was applied to both datasets to preserve EMG-EEG trial correspondence.

After preprocessing, the peak-to-peak MEP amplitude was computed per trial as the maximum-minus-minimum EMG within 20–50 ms post-stimulation. The mean MEP amplitude across trials was computed to index corticospinal excitability for each participant and condition. Participant-level iTEPs/TEPs were obtained by averaging EEG trials within each condition. To attenuate the ∼5 kHz TMS-pulse ringing artifact, TEPs were low-pass filtered using a time-domain Gaussian kernel (Figure S1). We computed the global mean field amplitude (GMFA) time course as a reference-free measure of global strength of EEG response. Participant-level iTEPs, TEPsand MEP traces across conditions are shown in Figures S2—S4.

To quantify iTEPs, we computed GMFA-AUC, defined as the area under the GMFA time course within predefined iTEP peak-aligned windows: 1.7–2.9 ms (Peak1), 2.9–4.4 ms (Peak2), and 4.4–6.0 ms (Peak3), consistent with our previous study [36]. We additionally quantified GMFA AUC in an early TEP window (6.0–22 ms) corresponding to TEP N15. TS-normalized inhibition indices were computed as paired pulse/TS ratios for each outcome (i.e., AUC_ratio = AUC_SICI/AUC_TS; MEP_ratio = MEP_SICI/MEP_TS), where values < 1 indicate inhibition relative to TS.

For the main experiment, positive and negative peaks were detected automatically from the iTEP trace at a channel close to the site of stimulation (channel 16 ) within predefined iTEP windows (1.7–6 ms) using MATLAB’s findpeaks function. For the follow-up dataset, positive peak 1 and peak 2 were searched. All detections were subsequently reviewed visually and manually corrected to eliminate false-positive detections.

### Statistical analysis

Statistical analyses were conducted in MATLAB and R (RStudio) on mean MEP amplitude, GMFA-AUC (peak1–3, N15), and TS-normalized inhibition indices (MEP_ratio; AUC_ratio), as defined above. Linear mixed-effects models were fitted in R (lme4). Fixed effects were tested using Type III Wald F-tests with Satterthwaite degrees of freedom (lmerTest). Unless otherwise stated, models were fitted using Restricted Maximum Likelihood (REML); nested model comparisons were conducted on maximum likelihood fits (REML = FALSE). Model assumptions (normality and homoscedasticity) were visually assessed using residual diagnostic plots. Where applicable, post-hoc pairwise contrasts were based on estimated marginal means with Holm-adjusted p-values.

Because hotspot targeting was optimized for FDI, primary EMG analyses were performed on the electromyographic signal recorded from the FDI muscles. Corresponding ADM results are summarized Figures S6 & S7. To test whether paired-pulse conditions modulated corticospinal output, we fitted mixed-effects models separately for FDI and ADM with condition (4 levels: TS/SICI_P1_/SICI_T1_/SICI_P2_) as a fixed effect and a random intercept for participant. Residual diagnostics indicated deviations from normality and heteroscedasticity on the original scale; therefore, participant-level mean MEP amplitude was converted to microvolt (V) and natural-log-transformed prior to model fitting:

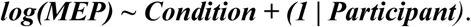

To evaluate condition effects on GMFA waveforms over time, we applied a nonparametric cluster-based permutation test [37]. Analyses were conducted in two predefined latency ranges (1.7–6 ms and 6–200 ms) , enabling separate assessment of early iTEP and later TEP activity and reducing the risk that larger later components obscure smaller early effects [38]. Statistical significance was evaluated under the null hypothesis that the GMFAs did not differ across four conditions (cluster threshold: p < .05; dependent F-test, critical α < 0.025; randomization = 10000). The critical α level was corrected for the two tested windows.

To make peak-specific inferences, GMFA-AUC in the iTEPs Peak1–3 windows was analyzed using a linear mixed-effects model with fixed effects of condition (4 levels: TS/SICI_P1_/SICI_T1_/SICI_P2_), peak (3 levels: Peak1–3), and their interaction, and a random intercept for participant:

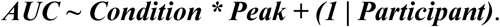

N15 AUC was analyzed using:

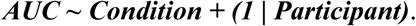

Because heteroscedasticity was indicated for N15 GMFA-AUC across conditions, N15 data was analyzed with a mixed-effects model allowing condition-specific residual variances. All inference and post-hoc contrasts were based on this variance-structured model.

To relate cortical and corticospinal inhibition, we tested associations between MEP_ratio and AUC_ratio for each AUC measure of interest. To test whether the AUC–MEP association differed by condition (3 levels: SICI_P1_/SICI_T1_/SICI_P2_), we compared an interaction model:

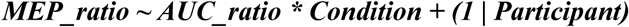

against an additive model:

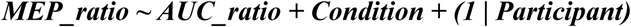

using likelihood-ratio tests on ML fits (REML = FALSE). If the interaction did not improve model fit, the additive model was refitted using REML for inference and reporting. The same procedure was applied for Peak2 and N15 predictors.

## Results

### Individual short-latency intracortical facilitation peaks and trough

During the experiment SICF curves were generated to establish the interstimulus intervals (ISI) that corresponded to the two facilitation maxima along with the interspersed minima (Figure 2). In all participants SICF displayed a bimodal distribution across the tested ISIs with minimal facilitation or even small inhibition as compared to the MEP amplitude evoked with single pulse TMS between 1.7 and 2.5ms (Figure 2A). The modes for the first peak, the first trough and the second peaks were 1.5ms, 2.1ms and 2.7ms (Figure 2B).

**Figure 2.**
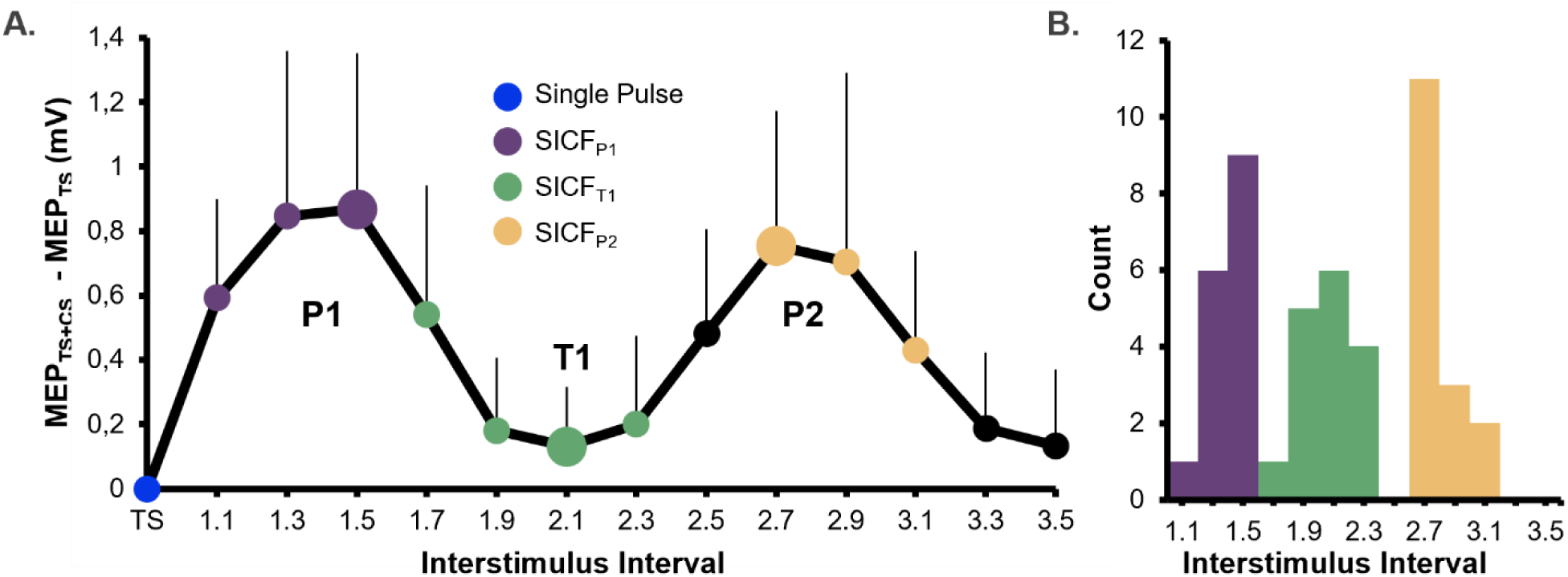
Measurements of short-latency intracortical facilitation (SICF). Single and paired biphasic stimuli were delivered over the left FDI hotspot and MEPs were recorded from the relaxed right FDI muscle. SICF was assessed at 13 paired-pulse intervals, ranging from 1.1 to 3.5 ms. **A.** Mean SICF curves (group level). The curve denotes the average difference between the median of paired-pulse amplitudes at a given ISI and median MEP response to the single-pulse test stimulus (TS). This method is adapted from Kesselheim et al., 2023 to avoid the large variability introduced when plotting response sizes relative to small single-pulse MEPs. The color of each dot represents if the corresponding ISI resulted in peak or trough facilitation for one or more subjects and large dots signify the mode of each peak and trough. The error bars denote standard deviation of the mean. **B.** Histogram over the distribution of ISIs corresponding to each peak and trough. To exemplify, neither peak 1 or 2 or the interspersed trough occurred at an ISI of 2.5 ms for any subjects, whereas the first peak facilitation (peak 1) was most frequently observed with and ISI of 1.5 ms. TS= Test Stimulus, CS= Conditioning Stimulus, P1=Peak 1, P2=Peak 2, T1 = Trough 1. *n* = 16.

### Individual short-latency intracortical inhibition (SICI) recruitment curves

#Figure 3 displays the relationship between CS intensity and MEP inhibition on the group level (Figure 3A) as well as the distribution of CS intensity (Figure 3B) corresponding to maximum MEP inhibition (Figure 3C). Maximal inhibition occurred at CS intensities between 65 and 105% of AMT with 95% AMT being the intensity that most frequently yielded the largest magnitude of inhibition. Individual maximal inhibition ranged from 92% to 29% with a mean of 68%.

**Figure 3.**
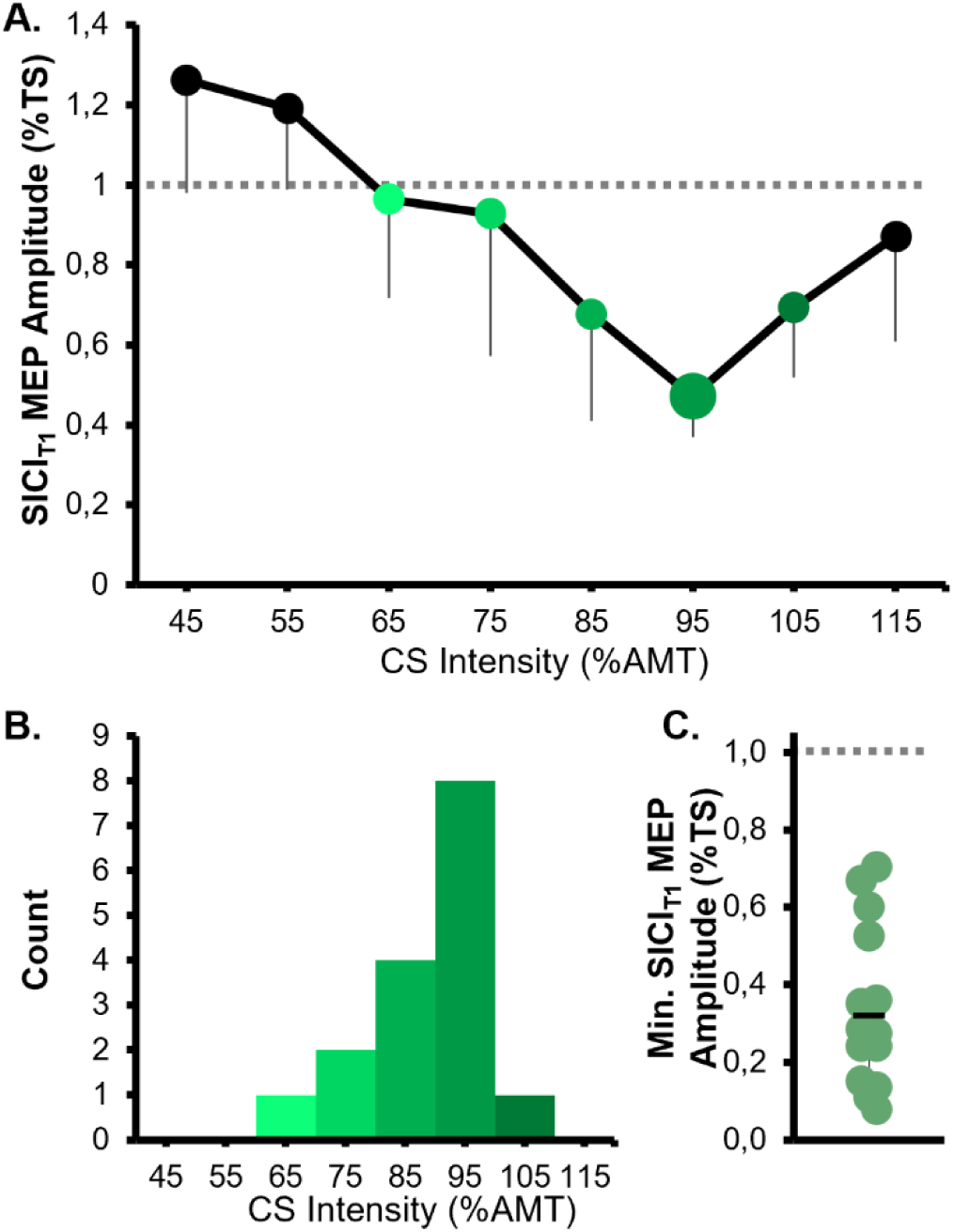
Establishing individual intensity for short-latency intracortical Inhibition (SICI). **A.** SICI recruitment curves representing the average paired pulse MEP amplitude relative to single-pulse MEP amplitude. Error bars are displayed unidirectional and represent Standard Error for visualization purposes. **B**. Histogram over the CS intensities that resulted in the largest MEP suppression. **C**. Individual median inhibition across 5 trials at the CS that caused peak MEP suppression indexed as the ratio between conditioned and test MEP. The black line signifies mean across 16 subjects and the grey dashed line corresponds to the test 100 percent of response amplitude. AMT = Active Motor Threshold, CS= Conditioning Stimulus, SICI= short-latency intracortical inhibition, T1= Trough 1, TS= Test Stimulus. *n* = 16.

### Effects of paired pulse on MEP amplitude in the main experiment

Compared with TS, the three paired-pulse SICI conditions (SICI_P1_, SICI_T1_, SICI_P2_) resulted in smaller MEP amplitudes (Figure 4A). This was supported by a significant main effect of condition on log-transformed MEP amplitude (F(3, 45) = 11.94, p < 0.0001). Post-hoc comparisons showed that each SICI condition produced smaller MEPs than the single-pulse TS (SICI_P1_ vs. TS: p = <.0.001; SICI_T1_ vs. TS: p = 0.001; SICI_P2_ vs. TS: p = 0.013). In addition, the shortest-ISI SICI condition yielded smaller MEPs than the longest-ISI condition (SICI_P1_ vs SICI_P2_: p = 0.026). Notably, MEP inhibition was not observed in all participants, as indicated by MEP_ratio ≥1 in Figure 4B.

**Figure 4.**
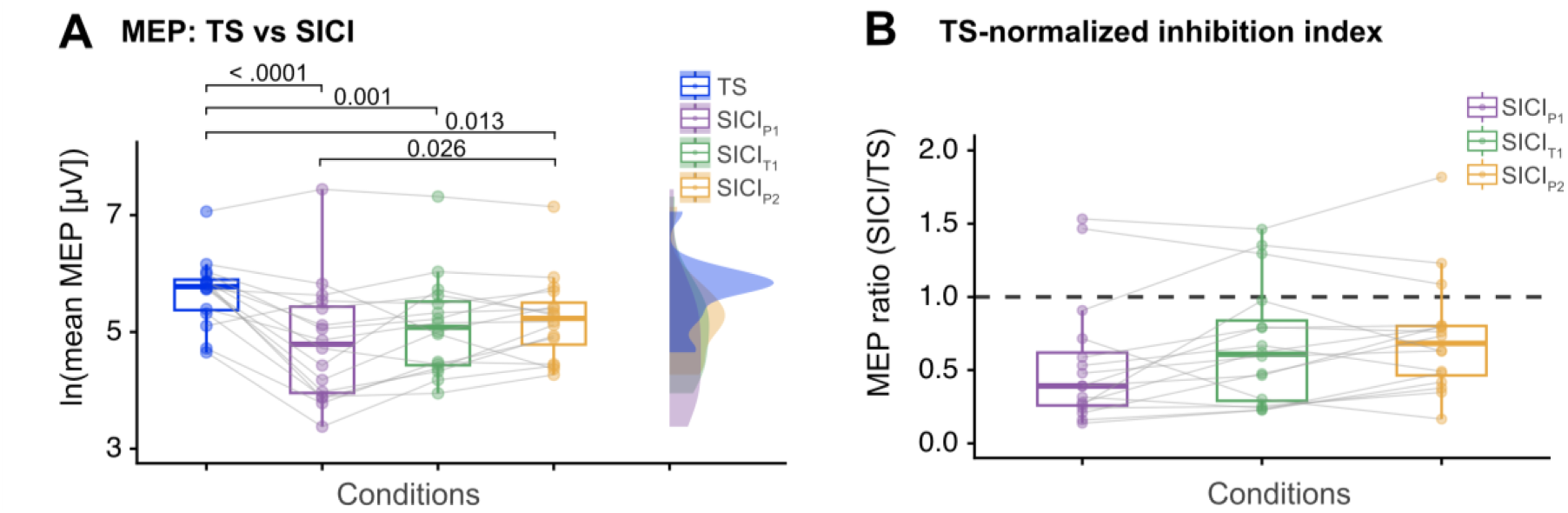
Paired-pulse MEP inhibition in the main experiment. **A.** Mean MEP amplitudes from the single-pulse condition (TS) and the three paired-pulse SICI conditions (SICIP1, SICIT1, SICIP2) amplitude in the targeted muscle FDI. Log-transformed values are shown as boxplots and density plots, with overlaid points for individual subjects (n=16) and grey lines indicating within-subject changes. ISIs in SICIP1, SICIT1, and SICIP2 were adjusted to Peak 1 (P1), Trough 1 (T1), and Peak 2 (P2) of the individual SICF curve, respectively. Significant differences in post-hoc tests are indicated with horizontal lines with Holm-adjusted p-values noted above. **B**. Relative MEP inhibition, expressed as the ratio of paired-pulse (SICIP1, SICIT1, and SICIP2) to single-pulse TS MEP amplitude, shown as boxplots with overlaid points for individual subjects and grey lines indicating within-subject changes. The dashed horizontal line at 1 marks the TS reference level. ISI = interstimulus interval, SICI = short-latency intracortical inhibition, SICF= short-latency intracortical facilitation, TS = Test Stimulus. *n* = 16.

### Effects of paired pulse on immediate TEPs in the main experiment

Single-pulse TMS elicited the characteristic iTEP waveform (1.7–6 ms) with three rapidly changing peaks superimposed on a slower component (Figure 5A), consistent with our previous work [30,31]. Across ISIs, SICI attenuated the iTEP response, with the most consistent suppression occurring around the second and third iTEP peaks. Peak latencies for each condition and subject (when peaks could be automatically detected) are provided in Table S3.

**Figure 5.**
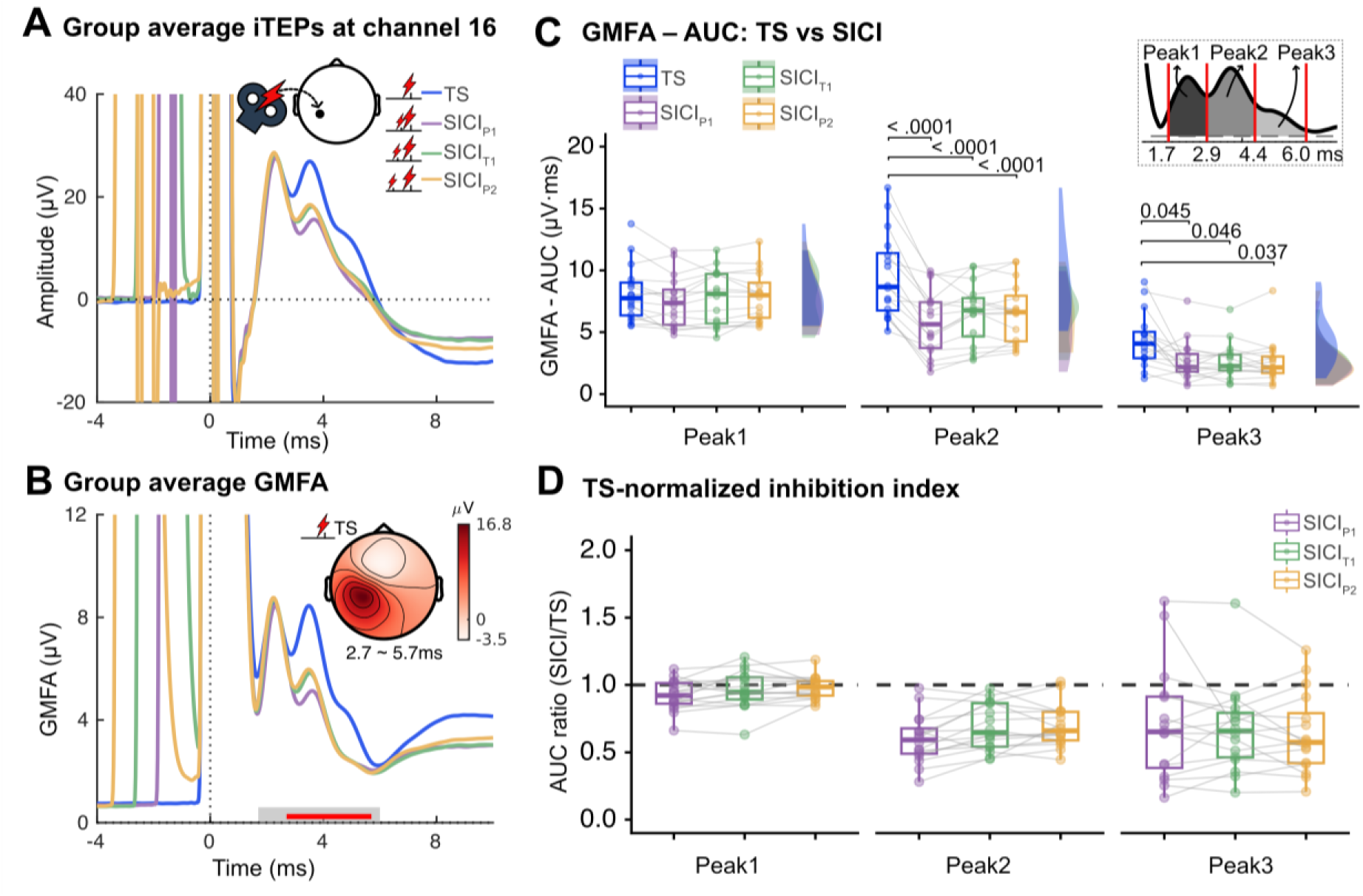
Paired-pulse iTEP inhibition in the main experiment. **A**. Group-average iTEPs at a channel close to the site of stimulation (channel 16) for the single-pulse condition (TS) and the three paired-pulse SICI conditions (SICIP1, SICIT1, SICIP2). ISIs in SICIP1, SICIT1, and SICIP2 were adjusted to Peak 1 (P1), Trough 1 (T1), and Peak 2 (P2) of the individual SICF profile, respectively. **B**. Group-average GMFA from the same conditions as in A. The thick grey shaded line represents the search window used for cluster-based permutation tests, and the red line indicates the temporal extent of the cluster driving the condition effect. The inset shows the scalp topography of the mean iTEP amplitude over 2.7–5.7 ms, corresponding to the identified cluster. **C.** Extracted AUC computed within predefined iTEPs peak-aligned windows: 1.7–2.9 ms (Peak1), 2.9–4.4 ms (Peak2), and 4.4–6.0 ms (Peak3), as shown in the inset. Significant differences in post-hoc tests are indicated with horizontal lines with Holm-adjusted p-values noted above. **D.** Relative iTEPs inhibition, expressed as the ratio of paired-pulse (SICIP1, SICIT1, and SICIP2) to single-pulse TS AUC value for the three time windows shown in C, displayed as boxplots with overlaid points for individual subjects and grey lines indicating within-subject changes. The dashed horizontal line at ‘1’ marks the TS reference level. AUC = Area under curve, ISI = interstimulus interval, GMFA= Global Mean Field Amplitude, SICI = short-latency intracortical inhibition, SICF= short-latency intracortical facilitation, TS = Test Stimulus. *n* = 16.

We statistically evaluated iTEP modulation using a two-step procedure. First, a cluster-based permutation test on the GMFA time course (1.7–6 ms) revealed a significant condition effect driven by a cluster spanning approximately 2.7–5.7 ms, overlapping the later iTEP components (Figure 5B). Second, to quantify peak-specific effects, we computed the AUC of the GMFA waveform within predefined iTEP peak-aligned windows (GMFA-AUC Peak1–3) as described in the Methods. A linear mixed-effects model showed a significant condition-by-peak interaction (F(6, 165) = 3.503, p = 0.003), indicating differential effects of SICI across iTEP peaks. Post-hoc contrasts demonstrated that, relative to TS, both GMFA-AUC Peak2 (all p < 0.0001) and Peak3 (all p < 0.05) were reduced for all three SICI ISIs, whereas Peak1 was not significantly affected (Figure 5C).TS-normalized iTEP inhibition indices are shown in Figure 5D. Upon visual inspection, there was a divergence between paired-pulse effect on global response strength indexed by GMFA, and on local iTEP morphology at electrodes near the stimulation site (Figure S5). This observation motivated a conservative focus on the more consistently suppressed Peak2 index in the coupling analyses below.

### Effects of paired pulse on later TEP in the main experiment

Later TEP components in the TS condition resembled the prototypical TEP morphology described in our previous work [31,39] (Figure 6A). For consistency with conventional nomenclature, we refer to the early negative deflection and its surrounding AUC window as N15, although it peaked slightly earlier than 15 ms in the present dataset, as in our prior work [26,27,39]. Across analyses, the most robust SICI-related effect was confined to this early window: a cluster-based permutation test on the GMFA time course (6–200 ms) identified a significant condition effect driven by a cluster spanning approximately 7–22 ms (Figure 6B), overlapping the predefined N15 window (6–22 ms). Consistent with this result, N15 GMFA-AUC analyzed using a mixed-effects model with a condition-specific residual variance structure (see Methods) showed a significant main effect of condition (χ²(3) = 12.11, p = 0.007). Post-hoc contrasts indicated reduced N15 AUC for the shortest-and medium-ISI SICI conditions relative to TS (SICI_P1_ vs. TS: p = 0.014; SICI_T1_ vs. TS: p = 0.015), whereas the longest-ISI condition did not differ reliably from TS (SICI_P2_ vs. TS: p = 0.182; Figure 6C). TS-normalized N15 inhibition indices are shown in Figure 6D.

**Figure 6.**
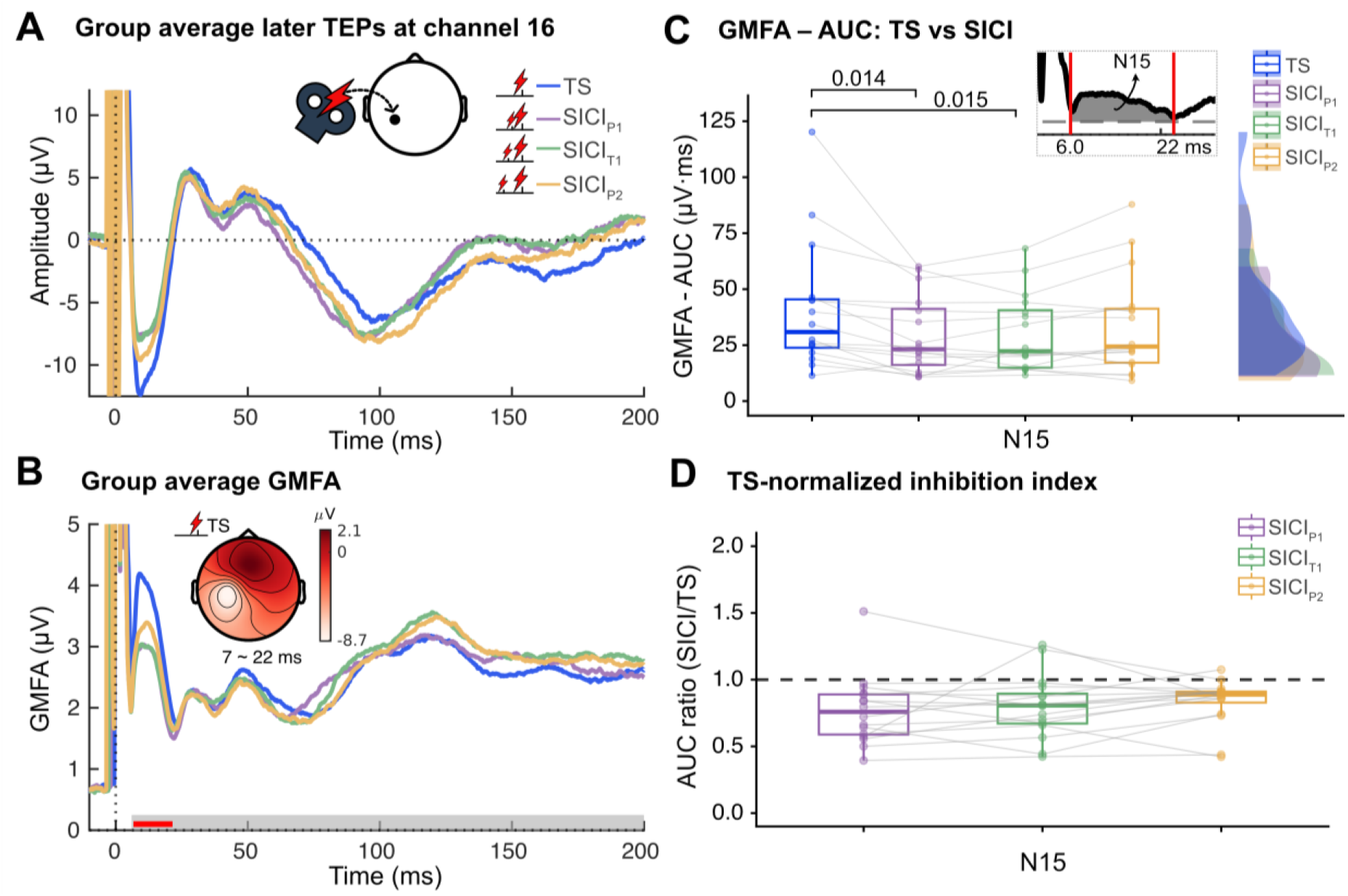
Paired-pulse later TEP inhibition in the main experiment. **A**. Group-average TEPs at EEG channel 16 near the stimulation site for the single-pulse condition (TS) and the three paired-pulse SICI conditions (SICIP1, SICIT1, SICIP2). ISIs in SICIP1, SICIT1, and SICIP2 were adjusted to Peak 1 (P1), Trough 1 (T1), and Peak 2 (P2) of the individual SICF curve, respectively. **B**. Group-average GMFA from the same conditions as in A. The thick grey shaded line represents the search window used for cluster-based permutation tests, and the red line indicates the temporal extent of the cluster driving the condition effect. The inset shows the scalp topography of the mean TEP amplitude over 7–22 ms, corresponding to the identified cluster. **C.** Extracted AUC computed within predefined N15 windows (6–22 ms), as shown in the inset. Significant differences in post-hoc tests are indicated with horizontal lines with Holm-adjusted p-values noted above. **D.** Relative TEP N15 inhibition, expressed as the ratio of paired-pulse (SICIP1, SICIT1, and SICIP2) to single-pulse TS AUC value for the time window shown in C, displayed as boxplots with overlaid points for individual subjects and grey lines indicating within-subject changes. The dashed horizontal line at ‘1’ marks the TS reference level. AUC = Area under curve, ISI = interstimulus interval, GMFA= Global Mean Field Amplitude, SICI = short-latency intracortical inhibition, SICF= short-latency intracortical facilitation, TS = Test Stimulus. *n* = 16.

### Associations between TEP inhibition and MEP inhibition in the main experiment

Cortical inhibition in the iTEP Peak2 window, but not N15, covaried with corticospinal inhibition, independently of SICI conditions. This was supported by linear mixed-effects regression analyses relating AUC_ratio to MEP_ratio. Based on the considerations on the robustness of the third iTEP peak described above, we focused on AUC_ratio for Peak2 and AUC_ratio for N15. For Peak2, likelihood-ratio testing provided no evidence that the AUC–MEP association differed across SICI ISIs (interaction vs additive model: χ²(2) = 1.948, p = 0.378). We therefore report the additive model, which showed a significant association between AUC_ratio for Peak2 and MEP_ratio (β = 0.863, p = 0.010; Figure 7A), indicating that greater cortical suppression in the Peak2 window coupled with greater corticospinal suppression. In contrast, AUC_ratio for N15 did not show a reliable association with MEP_ratio (β = 0.206, p = 0.335; Figure 7B).

**Figure 7.**
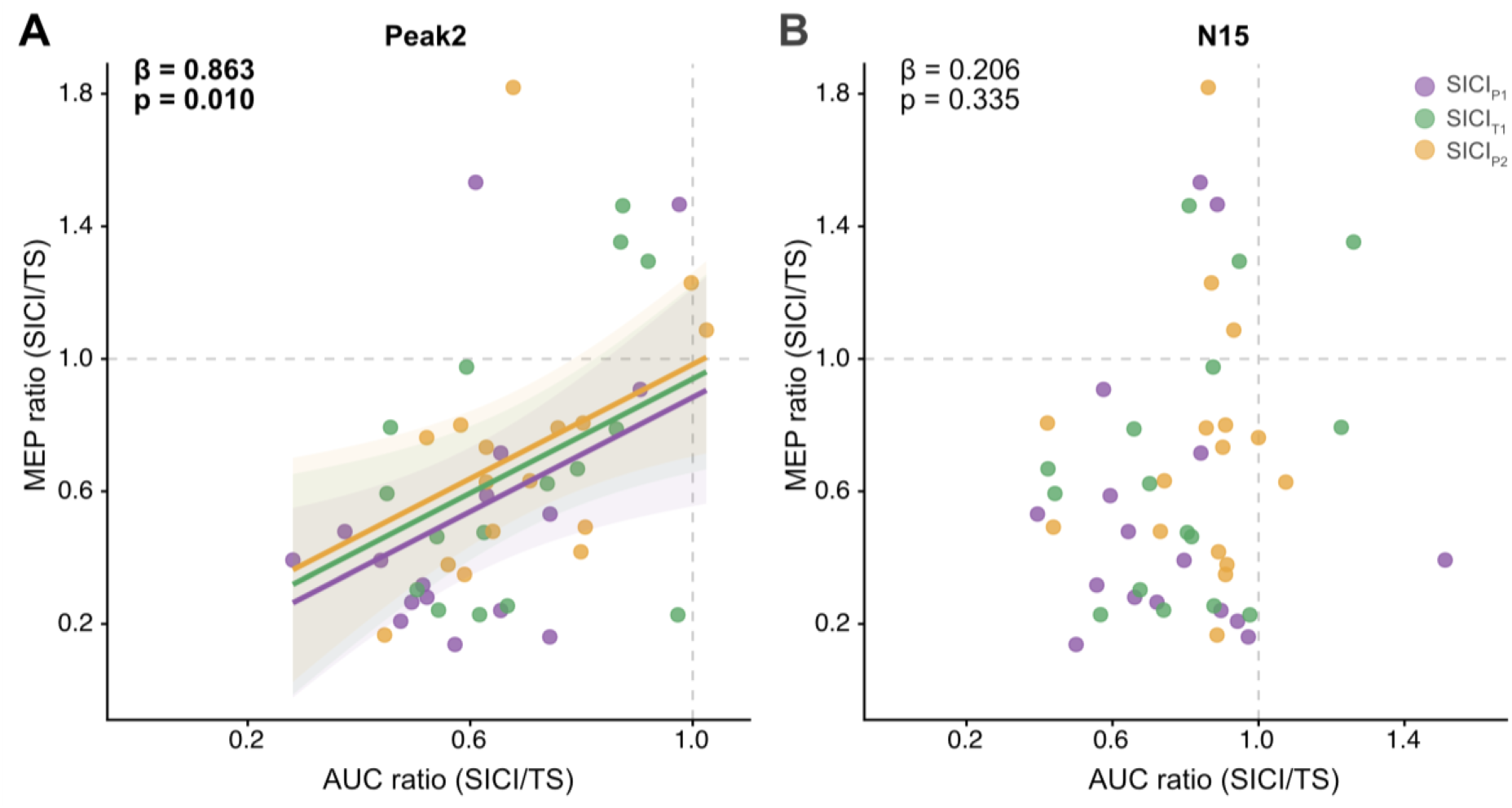
Relationship between cortical and corticomotor inhibition in the main experiment. **A**. Paired-pulse (SICIP1, SICIT1, and SICIP2) to single-pulse TS AUC ratio in the iTEP Peak2 time window, plotted against the corresponding paired-pulse to single-pulse MEP ratio. ISIs in SICIP1, SICIT1, and SICIP2 were adjusted to Peak 1 (P1), Trough 1 (T1), and Peak 2 (P2) of the individual SICF curve, respectively. The regression coefficient β and corresponding p-value are noted in the upper-left corner. A significant association, indicated by the fitted lines, was observed after adjusting for condition. **B**. Same as in A, except that the paired-pulse to single-pulse TS AUC ratio was computed for the TEP N15 time window. *n* = 16.

### Effect of CS intensity on iTEP, N15and MEP inhibition in the follow-up experiment

In a small cohort (*n*=4), we mapped the relationship between CS intensity and suppression of iTEPs, N15 and MEP. We found a consistent pattern of iTEP suppression well below the threshold for evoking MEP in the contracting FDI. Inhibition could be observed both when the CS intensity evoked a small iTEP peak and when it did not (see SUB2_004 vs SUB2_002 in Figure 8A). Although GMFA-AUC increased with stimulation intensity for both Peak1 and Peak2, distinct iTEP emergence at channel 16 was observed only for Peak1, indicating that Peak1 can be elicited at lower subthreshold intensities (Figure 8B). TS-normalized iTEP Peak2, TEP N15 and MEP inhibition indices are shown in Figure 8C. Peak2 AUC displayed a consistent pattern of inhibition with onset well below AMT_FDI_, and increasing with stimulation intensity. N15 AUC also expressed inhibition to varying degrees across the tested CS intensities, but with seemingly less consistency within and between-subject. MEPs displayed the strongest and most consistent suppression at the highest intensity (90% AMT), closely resembling the MEP SICI recruitment curve shown in Figure 3A

**Figure 8.**
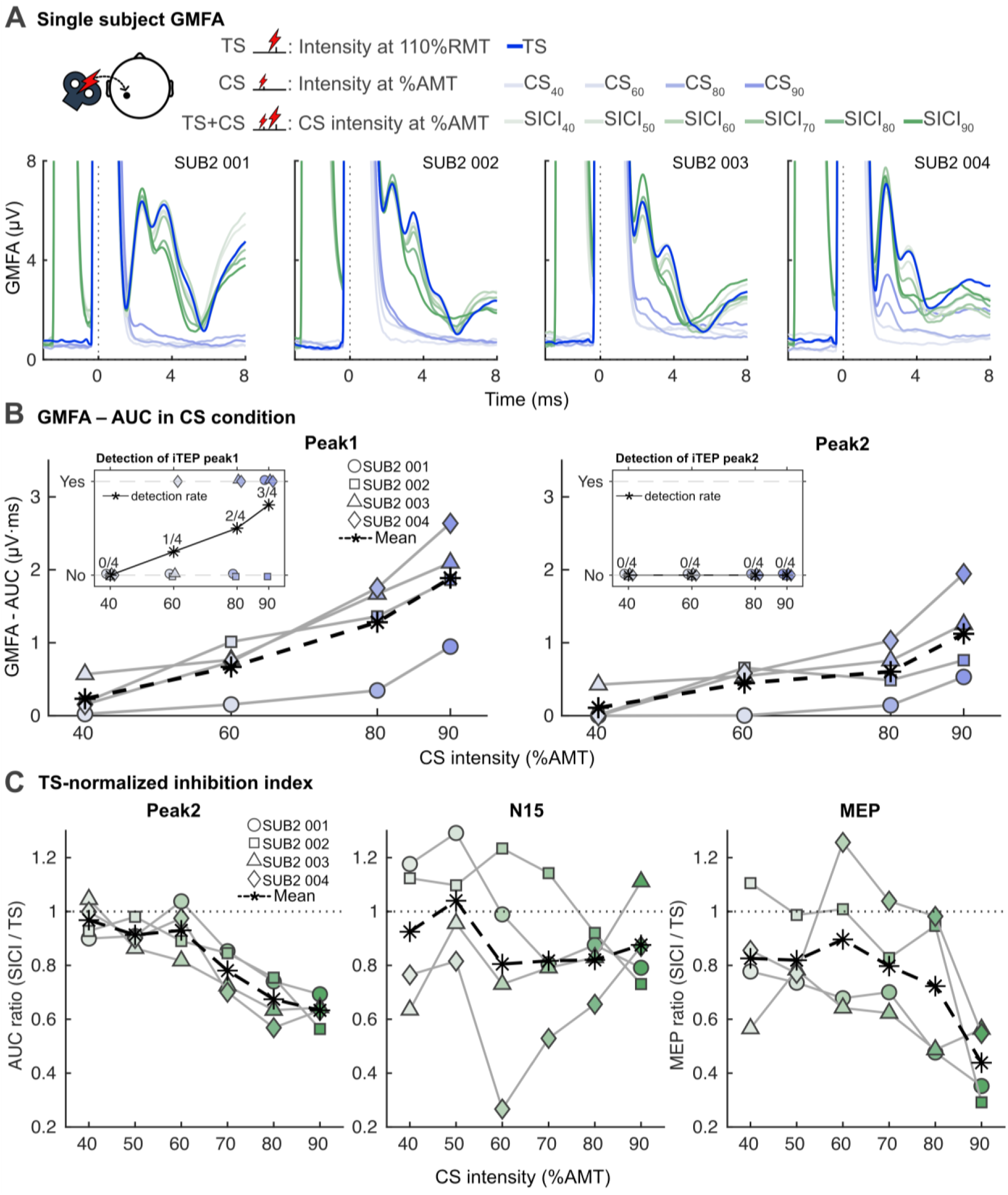
Relationship between CS intensity, iTEP and TEP inhibition. **A.** Individual GMFA traces from four subjects for single-pulse conditions (TS and CS; blue) and paired-pulse SICI conditions (green). CS intensity was set to 40%, 60%, 80%, and 90% AMT in CS40, CS60, CS80 and CS90, respectively, and set to 40%∼90% AMT in SICI40 –SICI90 conditions. Lighter shades denote lower CS intensities. TS intensity was set to 110% RMT. **B.** AUC extracted from predefined iTEPs peak-aligned windows: 1.7–2.9 ms (Peak1; left), 2.9–4.4 ms (Peak2; right) for CS conditions. The dashed black line is the mean across four subjects. Insets show the proportion of subjects in whom peaks can be automatically detected from the iTEP trace at a channel near the stimulation site (channel 16). **C.** Relative inhibition of iTEPs (left), N15 (middle) and MEPs (right) as a function of CS intensity. Inhibition shown as the ratio of paired-pulse to single-pulse TS AUC value (or MEP amplitude). The dashed black line indicates the mean across four subjects. The dashed grey horizontal line at ‘1’ marks the TS reference level. AMT = Active Motor Threshold, AUC = Area under curves, CS = Conditioning Stimulus, GMFA= Global Mean Field Amplitude, RMT = Resting Motor Threshold, SICI = short-latency intracortical inhibition, TS = Test Stimulus. *n* = 4.

## Discussion

Combining an optimized TMS-EEG approach with the paired-pulse SICI paradigm, we were able, for the first time, to directly trace regional inhibition in the human motor cortex, expressed as a paired-pulse reduction of immediate transcranial evoked potentials (iTEPs). Specifically, the second and third iTEP peaks were markedly reduced when a suprathreshold test stimulus was preceded by a subthreshold conditioning stimulus, irrespectively of whether the ISI was adjusted to the first peak, first trough, or second peak of the individual SICF curve. We further demonstrated a reduction of the N15 component which, unlike suppression of iTEP peak 2, did not covary with paired-pulse MEP inhibition. We interpret these data as showing that iTEPs index inhibition in circuit-specific intracortical pathways.

### Paired pulse TMS inhibits MEPs and immediate TEPs

In line with a large body of literature, paired-pulse TMS produced a robust inhibition of MEP amplitudes [3]. Our peak-wise analysis of iTEPs revealed selective suppression of later, but not the first iTEP peak. The iTEP complex consists of highly synchronized, fast peak components superimposed on a slower wave that reflects less synchronized local activity [26,40]. The periodicity of these fast components, their dependence on the direction of the indued cortical current, and their spatial confinement to pericentral sensorimotor cortices [30, 31] mirror key features of the corticospinal I-waves recorded epidurally at cervical or thoracic levels following motor cortex stimulation [14,15,41]. Whether the cortically generated iTEP peaks are the cortical counterparts of the descending volleys remains to be conclusively demonstrated.

The present findings strengthen this proposed link: Di Lazzaro and colleagues showed that a SICI paired-pulse protocol reduces later I-waves (i.e. I2, I3 and I4) while sparing the first I-wave, paralleling SICI-related inhibition of the MEP [5,25,42,43]. Peak-selective suppression of later components has also been demonstrated in peri-stimulus time histogram analyses of paired-pulse inhibition [44]. These observations are consistent with our finding that SICI selectively suppresses the second and third iTEP peaks, supporting the interpretation that these peaks index intracortical activity, contributing to later descending corticospinal volleys.

### Effect of interstimulus interval on short-latency inhibition of iTEPs and MEPs

To assess the potential impact of the timing of pulse pairs, we examined short-latency inhibition at three ISIs which were individually adjusted to participants first and second peak and first trough of their MEP SICF curve (SICI_P1_/SICI_T1_/SICI_P2_). The motor evoked responses showed the strongest SICI at the shortest ISI (SICI_P1_) at which SICI was significantly stronger than at the longest ISI (SICI_P2_). This finding aligns with previous epidural I-wave recordings showing the strongest inhibition of late I-wave volleys at the shortest interval (1.1 to 1.7ms) [5,25,42,43]. Prior MEP work also reported peak SICI at ISIs corresponding to the first SICF trough [24] and disinhibition-mediated increases in SICF in triple-pulse paradigms [25]. These interactions between SICI and SICF were not retrieved in the present study, most likely due to methodological differences. For instance, the neuronal circuits mediating SICI have lower thresholds than those mediating SICF and thus, the CS intensities employed in the present study might have been insufficient to excite neuronal elements involved in SICF.

Our iTEP recordings revealed comparable paired-pulse inhibition of later iTEP peaks at all three ISIs, providing no indication for stronger iTEP suppression at shorter intervals or differences in inhibition depending on SICF rhythmicity. This negative finding does not imply an absence of ISI dependent effects, as such interactions might only emerge at specific settings in the paired-pulse parameter space. Future studies employing different stimulus intensities or timings are warranted to explore whether and how interactions between SICI and SICF circuits tune the resulting iTEP modulation.

### Paired pulse effects on early TEP components

Covering a time window from 6 to 200 ms after the test pulse, exploratory GMFA analysis identified a significant cluster around N15 peak latency, showing a transient suppression during paired-pulse compared to single-pulse conditions. To our knowledge only seven previous studies have investigated short-interval paired-pulse effects on TEP peaks [45–51], three of which examined the N15 [47,50,51]. Two studies reported N15 suppression [47,51]while one study [50] observed increased in early contralateral (‘P13’) and ipsilateral (‘N18’) components. Our finding adds further evidence to a paired-pulse suppression of the early N15 component. In contrast, we did not detect significant clusters at other latencies, diverging from mixed reports of SICI effects on P30, N45, P60, N100, and P180 (reported in 2/5, 3/6, 4/4, 2/6, and 2/5 studies, respectively). Such heterogeneity likely reflects methodological differences both in acquisition, pre-processing and feature extraction [52]. A comprehensive, harmonized, and adequately powered effort to benchmark TEP methods is warranted [53].

Many TEP studies do not report the N15 component because of its proximity to TMS evoked instrumental and muscular artefacts, and its neural generators remain poorly understood [52]. Our experimental approach minimizes the risk of such confounds, and the topographical distribution of evoked activity confirms a pericentral origin rather than a muscular artefact at the rim of the scalp. Evidence suggests a pericentral origin with sources being located in the precentral gyrus [27,54]. While the N15 has been proposed to reflect sustained activity of cortical circuits in the stimulated motor cortex [55,56] (see e.g. [57] for discussion), the N15 latency is compatible with reverberatory cortico-cortical or cortico-subcortico-cortical activity. Pharmacological (midazolam-induced) and behavioral (tonic contraction) reductions of N15 [39,58], are consistent with a corticothalamic or other cortico-subcortical reverberating mechanism. Our replication of N15 suppression by an inhibitory paired-pulse paradigm accords with GABAergic inhibition of intratelencephalic and corticothalamic projection neurons, reducing both sustained local and cortico-subcortical responses.

### MEP inhibition relates to immediate but not early TEP inhibition

At the individual level, iTEP suppression covaried positively with MEP suppression. We restricted correlational analyses to peak-2 (SICI_P2_), because peak 1 was SICI-insensitive and peak 3 could not be extracted with sufficient precision for within-subject correlations. Given the physiological chain of events with iTEPs driving descending activity culminating in MEPs, SICI of the iTEPs is likely caused by the same GABAergic intracortical mechanisms probed with MEP or descending I-wave recordings. Notably, we observed occasional cases of modest peak 2 suppression of the iTEP without measurable MEP reduction (Figures 7 & 8). Inter-individual variation in suppression of later iTEP peaks, and by analogy later I-waves, may contribute. [18]. Alternatively, spinal or motoneuronal excitability fluctuations may have masked a MEP inhibition despite reduced corticospinal descending volleys. Crucially, the ability to index cortical inhibition free from spinal confounds opens avenues to chart the full temporal extent of GABAergic inhibition, which may markedly outlast the conventional 4–5 ms window [3,44]. The iTEP-MEP association supports the view that iTEPs capture the initial cascade of cortical excitation that ultimately summates at spinal motoneurons to generate MEPs [26,27,40].

In contrast to iTEPs, the magnitude of N15 suppression did not correlate with MEP inhibition. This negative finding is consistent with some [47,50] but not all [53] reports and adds to evidence that N15 results from effective re-excitation of the targeted cortex via a cortico-cortical and cortico-subcortical-cortical cascades which are partially dissociable from corticomotor output [31,39,62]

### SICI Input/Output curves

In four participants, we explored the relationship between the intensity of the conditioning stimulus and iTEP, MEP and N15 inhibition. First, and in correspondence with majority of the participant in main experiment, the threshold for evoking SICI was lower than the one for evoking descending volleys indexed either as iTEPs or indirectly as the AMT. In contrast to N15 and MEP suppression, the suppression of iTEP peak 2 followed a clear pattern across all four participants, suggesting that N15 and MEP suppression are subject to confounding effects somewhere in the signaling cascade. Third, and in agreement with previous published iTEP and I-wave data [36,59], our biphasic AP-PA recruited the first peak at lower intensities than the second peak in all four participants. Together, the results suggest that iTEPs can reliably index cortical inhibition using CS intensities solidly below the threshold for evoking descending activity, and that cortical inhibitory and excitatory dynamics can now be studied with subthreshold intensities and independent of downstream confounders.

### Methodological considerations

MEP inhibition was not observed in all participants at all ISIs during the TMS-EEG experiment (Figure 4B), despite individualizing CS intensity based on peaks and troughs from SICI recruitment curves. Several factors may account for this: SICI curves were acquired with a 5-s inter-trial interval, whereas combined TMS-EEG/EMG recordings paused for two seconds between trials. While SICI appears stable at a 4-s inter-trial interval [60], effects at 2 s remain untested. Conversely, single-pulse MEPs at a 2-s inter-trial interval are smaller than at 10 s [61], potentially reducing the dynamic range for detecting inhibition and even producing apparent facilitation in some participant. Slow excitability drifts also need to be considered: Low-frequency single/paired-pulse stimulation over prolonged recordings (ITI = 2 s) may cause within-block drifts in corticospinal excitability. Although block order was randomized and counterbalanced (limiting group-level bias), selective within-block effects on paired vs single pulses cannot be excluded. Despite absent SICI in three participants, group-level effects were robust, and the increased within-subject variance in relative inhibition enabled informative correlations with immediate and early cortical markers.

We quantified inhibition of the first three iTEP peaks. A fourth peak is occasionally present but coincides with the descending slope of the N15 peak, impeding reliable extraction of peak 4 amplitude or area. Methods that separate fast (peak) and slow (envelope) components may expose additional inhibitory effects at later peaks. Likewise, area-under-the-curve (AUC) reductions cannot be uniquely attributed to peak vs slow-wave components. As illustrated in Figures 5A, 8A, S5, SICI appears to shorten the slow-wave duration, implying effects beyond elements generating the largest corticomotoneuronal volleys. (see [62]). Future work should develop decomposition approaches that preserve peak-wise quantification while isolating dynamics of the slow iTEP component.

Finally, in this and our previous experiments, we have dosed stimulation based on MEP thresholds. As the TEP optimized hotspot is generally not identical with the MEP hotspot, it warrants a cautious interpretation of relative intensity needed to evoke an iTEP signal or iTEP suppression. To clarify the actual activation and suppression thresholds, future studies can benefit from dosing relative to evoked electric fields or the muscle unspecific iTEP signal rather than the muscle specific MEP (see e.g. [63]).

## Conclusions and perspectives

The combination of a well-established short-latency paired-pulse paradigm with iTEP recordings yields a cortical read-out of SICI (i) that is expressed by the later iTEP components (i.e., iTEP peak 2 and peak 3), (ii) correlates with MEP suppression, and (iii) dissociates from the early N15 dynamics. Together, the findings support the view that SICI predominantly targets intracortical circuits generating later descending volleys, while N15 reflects parallel or downstream pericentral/cortico-subcortical processes.

An advantage of the short-latency paired-pulse TEP approach is that it is caused by cortical activity in response to TMS and therefore offers a more direct measure of TMS-induced cortical inhibition than MEPs, which are influenced by downstream transmission along the corticomotor pathway. Both, the iTEPs and the early N15 component exhibited paired-pulse inhibition, yet the magnitude and temporal profile of inhibition differed between these responses. This dissociation indicates that TMS engages at least two distinct cortical circuits: one underlying the generation of iTEPs and closely linked to corticomotor output, and another contributing to the N15 component. Together, these findings demonstrate that short-latency paired-pulse TEPs can dissociate functionally distinct inhibitory cortical mechanisms that are not resolvable using conventional MEP-based measures.

Future studies should (a) systematically vary ISIs and CS intensities to map threshold-dependent recruitment of SICI vs SICF, (b) incorporate source-localized TMS-EEG approaches to resolve generators of iTEP vs N15, (c) apply component-separation methods to disentangle fast peaks from the slow envelope, and (d) integrate epidural I-wave recordings where feasible to directly study the coupling between cortical iTEPs and corticospinal descending I-waves.

## Acknowledgements

The authors would like to thank Marten Nuyts and Agata Banach for their help in setting up the experiment, and Jonas Laugesen for assistance with data acquisition.

## Author contributions

**LC, KM, JR & HR** conceptualized and designed the study. **LC, YS, DH, MMB & KM** collected data. **LC, YS & DH** analyzed data. **LC & YS** drafted the first version of manuscript. **All authors** have critically reviewed the manuscript and approved of the final version.

