## Supplementary figures and tables for "Direct Assessment of Short-Latency Intracortical Inhibition via Immediate TMS-Evoked Potentials"

* Shared first authorship

### Supplementary Methods and Results

In the following sections, we provide additional methodological details and results that support our main conclusions.

#### Overview of number of trials and EEG channels for analysis

**Table S1** Overview of the experimental conditions, for each subject, the numbers of trials and EEG channels included for analysis. Note: due to technical errors, only 50 trials were recorded for SUB_015 (TS and CS) and SUB_016 (TS).

| | **TS** | | **$SICI_{P1}$** | | **$SICI_{T1}$** | | **$SICI_{P2}$** | | **CS** | |
|---|---|---|---|---|---|---|---|---|---|---|
| **SUBJECT** | **Trials** | **Channels** | **Trials** | **Channels** | **Trials** | **Channels** | **Trials** | **Channels** | **Trials** | **Channels** |
| SUB_001 | 65/100 | 52/61 | 50/100 | 54/61 | 72/100 | 54/61 | 61/100 | 54/61 | 60/100 | 53/61 |
| SUB_002 | 88/100 | 61/61 | 81/98 | 59/61 | 88/100 | 60/61 | 80/100 | 61/61 | 83/100 | 58/61 |
| SUB_003 | 52/100 | 51/61 | 62/100 | 50/61 | 83/100 | 50/61 | 57/100 | 50/61 | 61/100 | 49/61 |
| SUB_004 | 90/100 | 56/61 | 89/100 | 56/61 | 92/100 | 56/61 | 86/100 | 56/61 | 90/100 | 57/61 |
| SUB_005 | 74/100 | 57/61 | 86/100 | 56/61 | 69/100 | 56/61 | 85/100 | 57/61 | 73/100 | 56/61 |
| SUB_006 | 82/100 | 57/61 | 81/100 | 57/61 | 78/100 | 57/61 | 87/100 | 56/61 | 71/100 | 57/61 |
| SUB_007 | 82/100 | 53/61 | 74/100 | 52/61 | 76/100 | 52/61 | 70/100 | 52/61 | 67/100 | 52/61 |
| SUB_008 | 91/100 | 56/61 | 82/100 | 54/61 | 48/100 | 55/61 | 91/100 | 55/61 | 84/100 | 55/61 |
| SUB_009 | 69/100 | 55/61 | 78/97 | 55/61 | 73/100 | 54/61 | 81/100 | 54/61 | 83/100 | 56/61 |
| SUB_010 | 61/100 | 60/61 | 63/100 | 60/61 | 61/100 | 60/61 | 53/100 | 59/61 | 66/100 | 58/61 |
| SUB_011 | 95/100 | 55/61 | 89/100 | 55/61 | 74/100 | 55/61 | 83/100 | 55/61 | 86/100 | 55/61 |
| SUB_012 | 74/100 | 54/61 | 78/100 | 53/61 | 82/100 | 52/61 | 77/100 | 53/61 | 81/100 | 53/61 |
| SUB_013 | 43/100 | 59/61 | 59/100 | 59/61 | 51/100 | 59/61 | 45/100 | 59/61 | 51/100 | 59/61 |
| SUB_014 | 63/100 | 50/61 | 59/110 | 50/61 | 62/100 | 50/61 | 60/100 | 50/61 | 71/100 | 50/61 |
| SUB_015 | 43/**50** | 59/61 | 87/100 | 60/61 | 89/100 | 60/61 | 88/100 | 60/61 | 41/**50** | 59/61 |
| SUB_016 | 46/**50** | 58/61 | 87/100 | 57/61 | 85/100 | 57/61 | 83/100 | 57/61 | 75/100 | 57/61 |
| **Median** | **72/100** | **56/61** | **80/100** | **56/61** | **75/100** | **56/61** | **81/100** | **56/61** | **72/100** | **56/61** |

**Overview of participants and stimulation parameters from the experimental conditions**
**Table S2** Overview of participants and the parameters established during the experiments. Red numbers denote TS intensities that deviated from 110 % RMT. Intensities was either increased to ensure suprathreshold stimulation (resting MEPs in >50 % of trials, n=5) or decreased to avoid a scalp artefact (n=1).

| | | | THRESHOLDS | | INTERSTIMULUS INTERVALS | | | INTENSITIES | | |
|---|---|---|---|---|---|---|---|---|---|---|
| ID | AGE | SEX | RMT TEP Hotspot | AMT TEP Hotspot | P1 | T1 | P2 | TS (%MSO) | CS (%AMT) | CS (%MSO) |
| SUB 001 | 35 | M | 71 | 43 | 1,1 | 1,9 | 2,7 | 78 | 85 | 37 |
| SUB 002 | 27 | F | 79 | 48 | 1,3 | 1,9 | 2,7 | 93 | 95 | 46 |
| SUB 003 | 26 | F | 90 | 58 | 1,5 | 2,1 | 2,7 | 100 | 105 | 61 |
| SUB 004 | 22 | M | 73 | 50 | 1,5 | 2,3 | 2,7 | 80 | 95 | 48 |
| SUB 005 | 23 | F | 70 | 50 | 1,5 | 1,9 | 2,7 | 77 | 85 | 43 |
| SUB 006 | 26 | F | 61 | 41 | 1,3 | 1,7 | 2,7 | 67 | 95 | 38 |
| SUB 007 | 28 | F | 75 | 55 | 1,3 | 2,1 | 2,7 | 83 | 95 | 75 |
| SUB 008 | 30 | F | 50 | 42 | 1,5 | 2,1 | 2,7 | 60 | 85 | 36 |
| SUB 009 | 20 | F | 59 | 43 | 1,5 | 2,3 | 3,1 | 65 | 75 | 32 |
| SUB 010 | 27 | M | 60 | 45 | 1,5 | 2,1 | 2,7 | 66 | 65 | 29 |
| SUB 011 | 25 | M | 61 | 41 | 1,3 | 1,9 | 2,7 | 67 | 95 | 39 |
| SUB 012 | 25 | F | 75 | 63 | 1,5 | 2,1 | 2,9 | 83 | 95 | 60 |
| SUB 013 | 30 | M | 60 | 38 | 1,3 | 2,1 | 2,7 | 70 | 95 | 36 |
| SUB 014 | 26 | M | 75 | 52 | 1,5 | 2,3 | 2,9 | 83 | 95 | 49 |
| SUB 015 | 24 | M | 88 | 72 | 1,3 | 2,3 | 3,1 | 97 | 75 | 65 |
| SUB 016 | 27 | F | 75 | 57 | 1,5 | 1,9 | 2,9 | 87 | 85 | 46 |

**Effects of low-pass filters (LPF) on iTEPs**

All three tested LPFs produced adequate attenuation at ~5 kHz, the TMS-pulse ringing artifact (Figure S1). The gaussian-kernel filter exhibited the least ripple in the impulse response compared to the Butterworth and windowed-sinc filters. Among them, the resulting iTEPs were largely similar in overall amplitude, with the gaussian filter introducing the smallest 'ripple' in the time domain waveform (Figure S1 C).

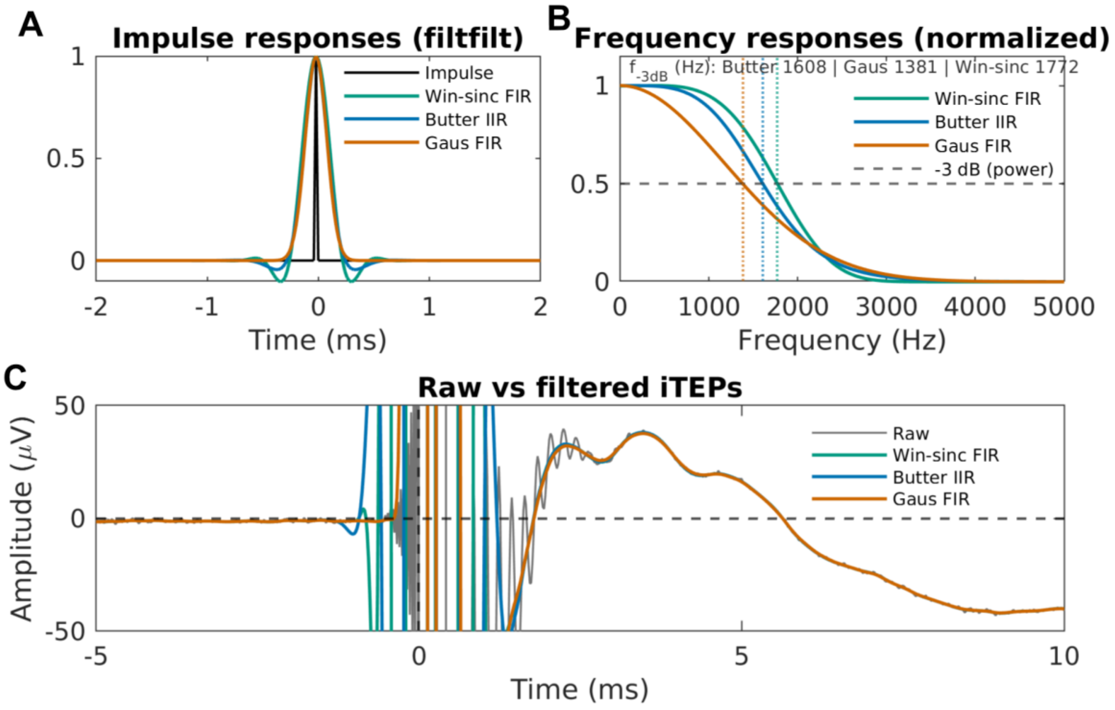


**Figure S1.** Comparison of three zero-phase low-pass filters (LPF) applied to iTEPs to attenuate TMS-pulse ringing (~5 kHz). Three filters are a 2nd-order Butterworth LPF (butter, cutoff 2 kHz), a time-domain Gaussian kernel LPF (FWHM = 0.16 ms; 25 samples), and a windowed-sinc LPF (fir1, order = 50, cutoff = 2.5 kHz), all applied with matlab function filtfilt. Pannels show the effective impulse responses and normalized frequency response of each filter, and the corresponding filtered iTEP waveforms from a single subject (TS condition). FWHM = Full Width Half Maximum, TS = Test Stimulus.

**Single-subject iTEP and N15 TEP waveforms, corresponding GMFA, and mean MEP traces across condition**

Across all subjects, iTEPs were consistently observed at the TS and the three paired-pulse conditions (Figure S2). In contrast, iTEPs from the CS condition were less clearly expressed, lacking high frequency i-TEP peaks. Notably, SUB_012 showed a clear iTEPs at CS condition but with markedly small amplitude compared to other conditions. Figure S3 provides a zoomed view of the TEP N15 time window. No prominent TMS-evoked muscle artifacts are visible, supporting the adequacy of preprocessing in this interval. Mean MEP trace for each subject is shown in Figure S4.

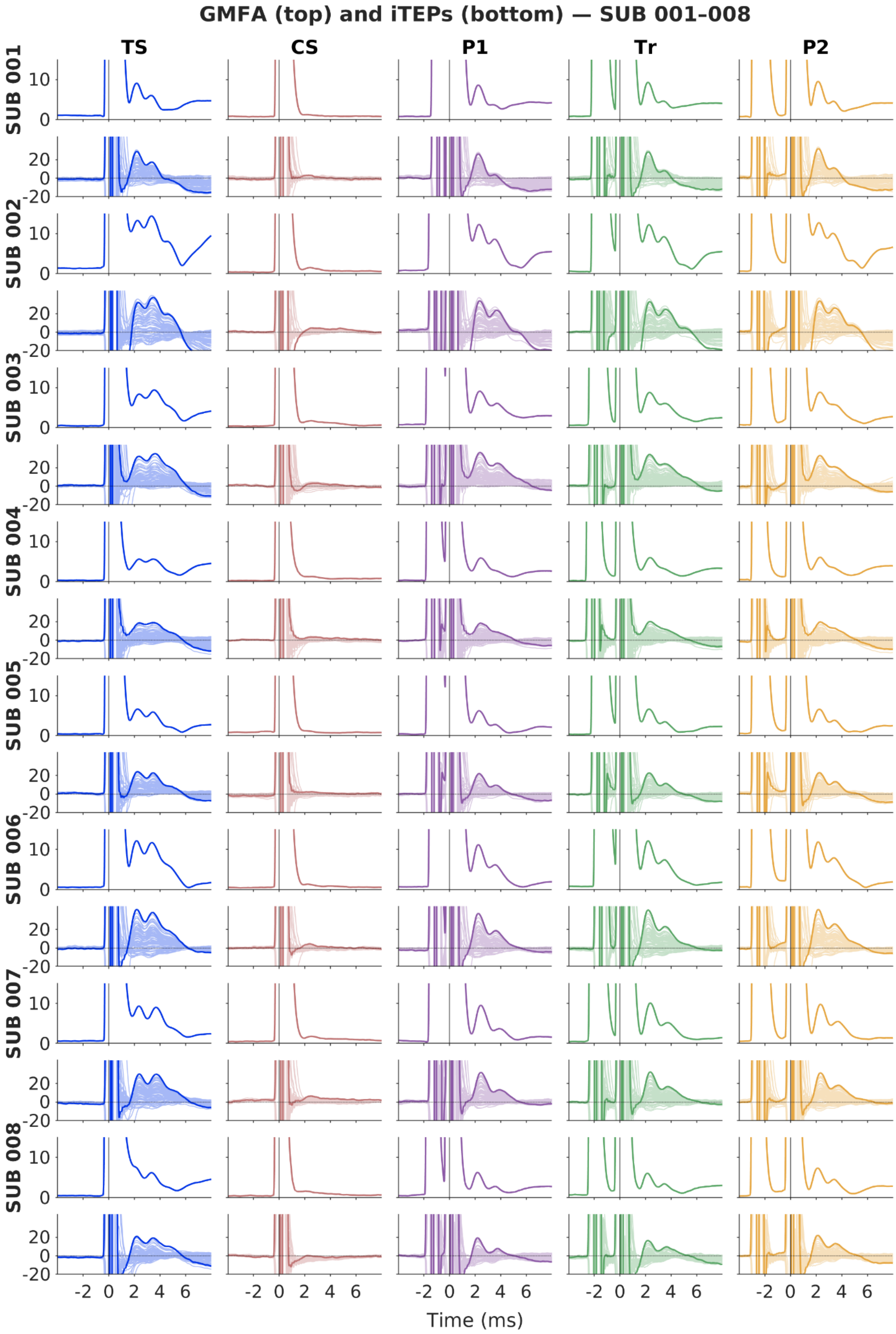

GMFA (top) and iTEPs (bottom) — SUB 001-008
TS
CS
P1
Tr
P2
SUB 001
SUB 002
SUB 003
SUB 004
SUB 005
SUB 006
SUB 007
SUB 008
Time (ms)

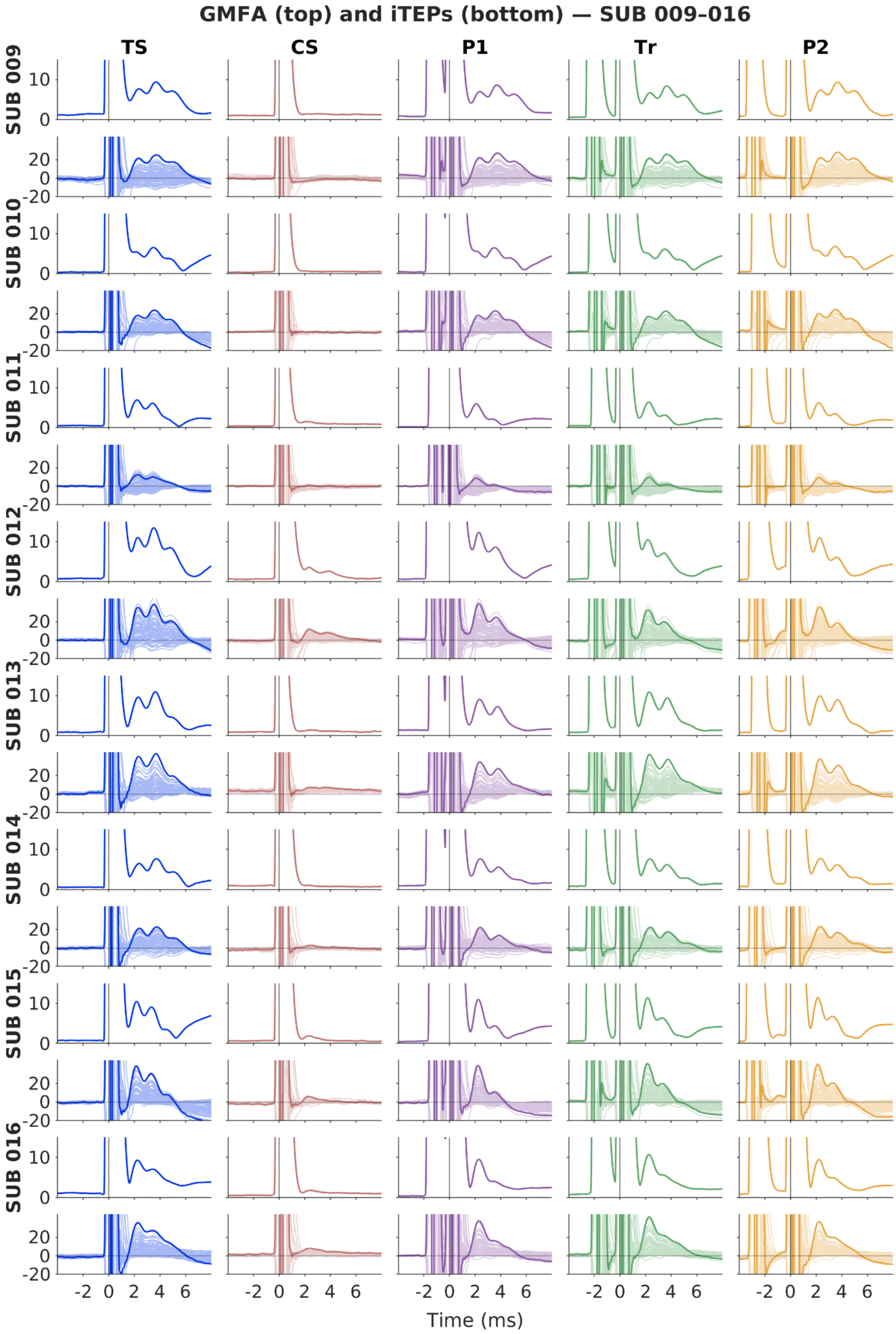
GMFA (top) and iTEPs (bottom) — SUB 009-016
TS
CS
P1
Tr
P2
SUB 009
SUB 010
SUB 011
SUB 012
SUB 013
SUB 014
SUB 015
SUB 016
10
0
20
0
-20
-2 0 2 4 6
Time (ms)

**Figure S2.** Single-subject **iTEP** at a channel near the site of stimulation (channel 16) and GMFA for the single-pulse conditions (TS and CS) and the three paired-pulse SICI conditions ($SICI_{P1}$, $SICI_{T1}$, $SICI_{P2}$). ISIs in $SICI_{P1}$, $SICI_{T1}$, and $SICI_{P2}$ were adjusted to Peak 1 (P1), Trough 1 (T1), and Peak 2 (P2) of the individual SICF curve, respectively.
CS = Conditioning Stimulus, ISI = interstimulus interval, GMFA= Global Mean Field Amplitude, SICI = short-latency intracortical inhibition, SICF= short-latency intracortical facilitation, TS = Test Stimulus.

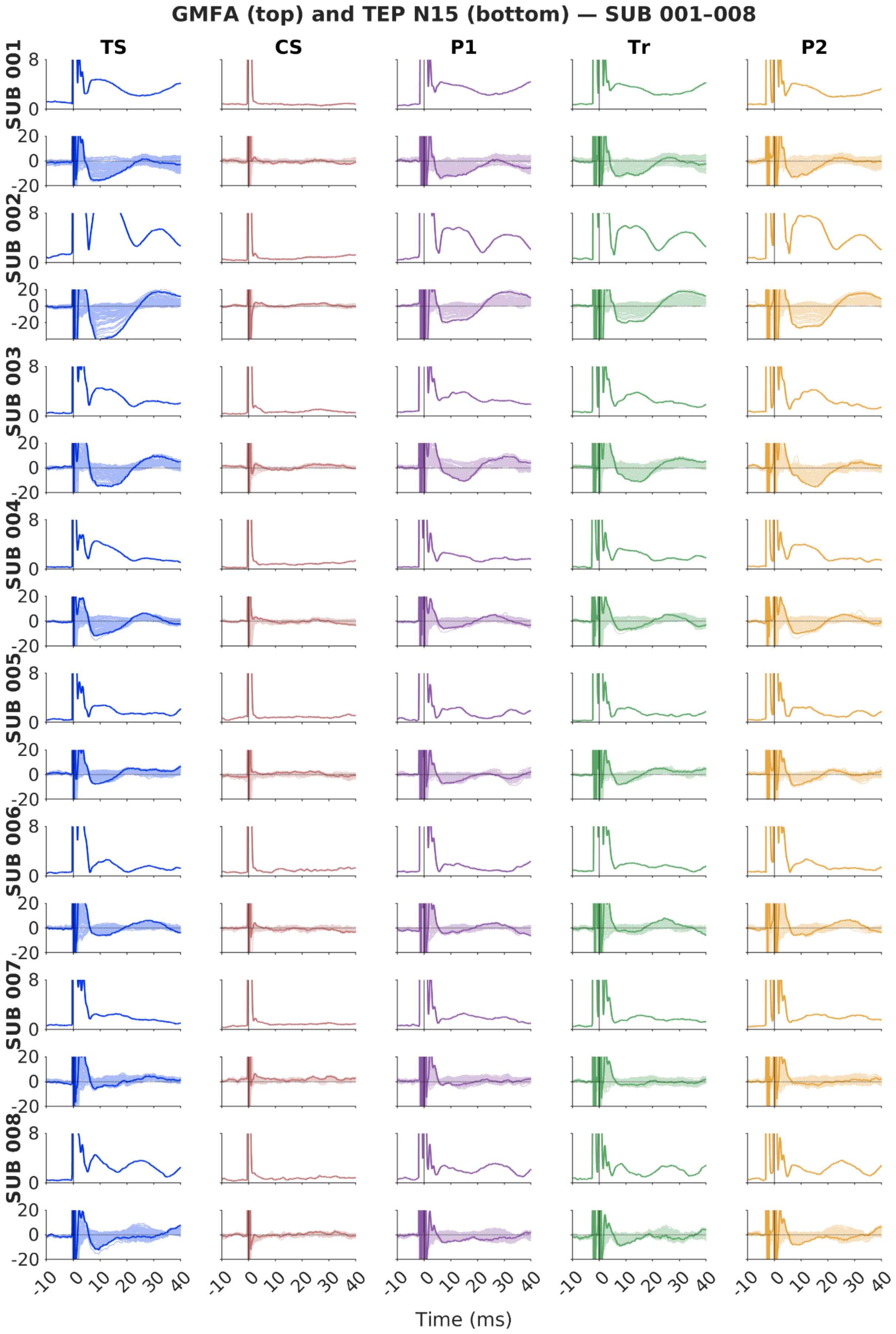
GMFA (top) and TEP N15 (bottom) — SUB 001–008
TS
CS
P1
Tr
P2
SUB 001
SUB 002
SUB 003
SUB 004
SUB 005
SUB 006
SUB 007
SUB 008
8
0
20
0
-20
-10
0
10
20
30
40
Time (ms)

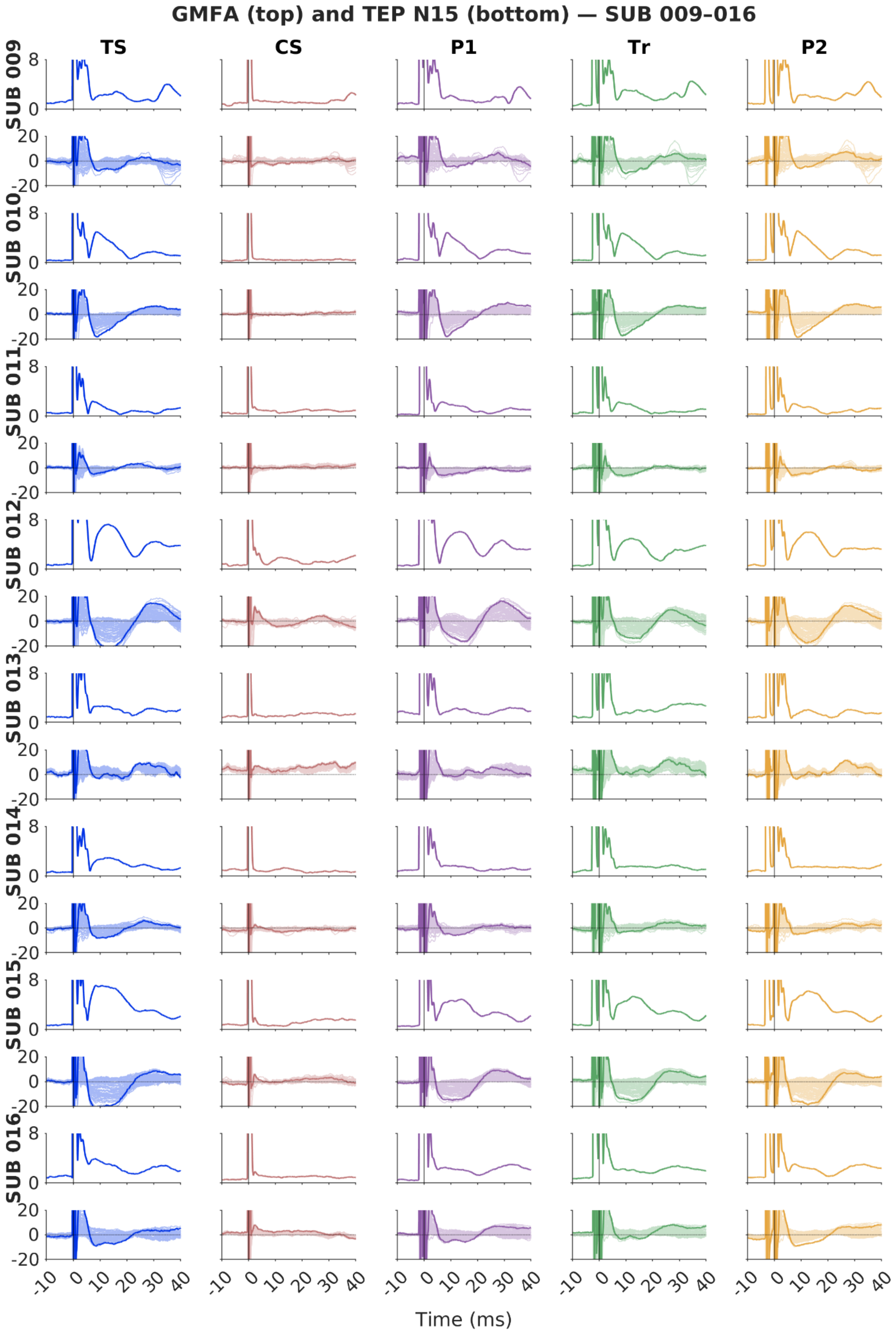

GMFA (top) and TEP N15 (bottom) — SUB 009–016
TS
CS
P1
Tr
P2
SUB 009
SUB 010
SUB 011
SUB 012
SUB 013
SUB 014
SUB 015
SUB 016
8
0
20
0
-20
-10
0
10
20
30
40
Time (ms)

**Figure S3.** Single-subject **TEP N15** at a channel near the site of stimulation (channel 16) and GMFA for the single-pulse conditions (TS and CS) and the three paired-pulse SICI conditions ($SICI_{P1}$, $SICI_{T1}$, $SICI_{P2}$). ISIs in $SICI_{P1}$, $SICI_{T1}$, and $SICI_{P2}$ were adjusted to Peak 1 (P1), Trough 1 (T1), and Peak 2 (P2) of the individual SICF curve, respectively.
CS = Conditioning Stimulus, ISI = interstimulus interval, GMFA= Global Mean Field Amplitude, SICI = short-latency intracortical inhibition, SICF= short-latency intracortical facilitation, TS = Test Stimulus.

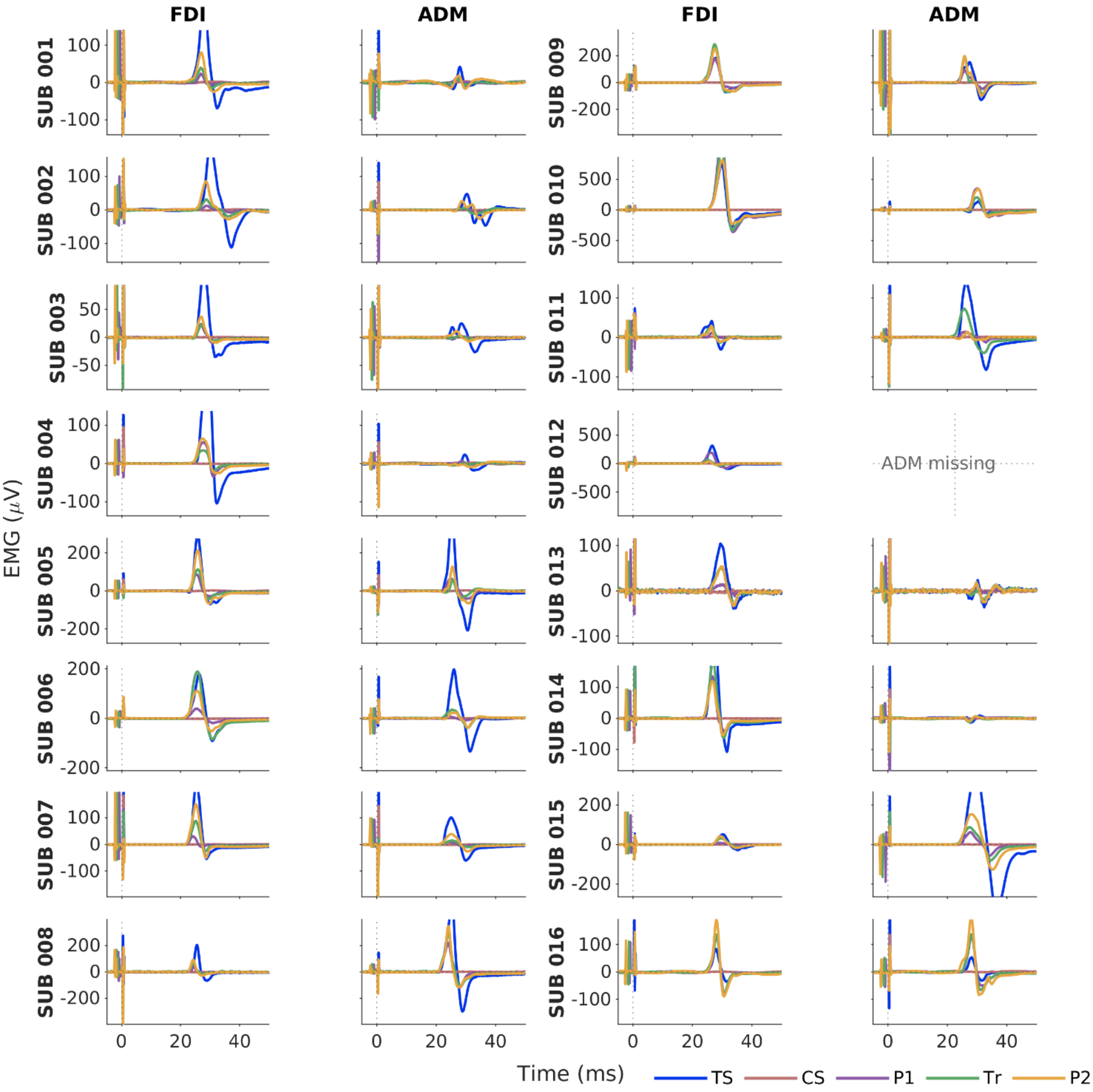


**Figure S4.** Single-subject mean MEP traces in hand muscles FDI and ADM for the single-pulse conditions (TS and CS) and the three paired-pulse SICI conditions ($SICI_{P1}$, $SICI_{T1}$, $SICI_{P2}$). ISIs in $SICI_{P1}$, $SICI_{T1}$, and $SICI_{P2}$ were adjusted to Peak 1 (P1), Trough 1 (T1), and Peak 2 (P2) of the individual SICF curve, respectively.
CS = Conditioning Stimulus, ISI = interstimulus interval, SICI = short-latency intracortical inhibition, SICF= short-latency intracortical facilitation, TS = Test Stimulus.

**Overview of peak and trough latencies and amplitudes from the iTEP waveform at an electrode near the stimulation site (channel 16)**

**Table S3.1** Peak/trough latency (ms) and amplitude (μV) for iTEP by participant at **TS condition**

| SUBJECT | Peak 1 | | Trough 1 | | Peak 2 | | Trough 2 | | Peak 3 | |
|---|---|---|---|---|---|---|---|---|---|---|
| SUBJECT | Lat (ms) | Amp (μV) | Lat (ms) | Amp (μV) | Lat (ms) | Amp (μV) | Lat (ms) | Amp (μV) | Lat (ms) | Amp (μV) |
| SUB_001 | 2.2 | 28.83 | 2.9 | 14.26 | 3.4 | 17.51 | 4.5 | -0.95 | NaN | NaN |
| SUB_002 | 2.3 | 32.09 | 2.8 | 25.59 | 3.5 | 37.59 | 4.4 | 19.20 | 4.6 | 19.65 |
| SUB_003 | 2.3 | 32.23 | 3.0 | 27.45 | 3.6 | 35.22 | NaN | NaN | NaN | NaN |
| SUB_004 | 2.4 | 18.71 | 2.9 | 17.14 | 3.5 | 19.11 | NaN | NaN | NaN | NaN |
| SUB_005 | 2.3 | 24.12 | 3.0 | 17.00 | 3.5 | 22.78 | NaN | NaN | NaN | NaN |
| SUB_006 | 2.2 | 41.49 | 2.8 | 28.27 | 3.4 | 38.38 | NaN | NaN | NaN | NaN |
| SUB_007 | 2.4 | 30.17 | 3.0 | 19.44 | 3.7 | 29.96 | NaN | NaN | NaN | NaN |
| SUB_008 | 2.2 | 21.12 | 2.9 | 13.48 | 3.5 | 19.35 | 4.6 | 10.22 | NaN | NaN |
| SUB_009 | 2.3 | 21.69 | 3.0 | 16.24 | 3.8 | 25.07 | 4.7 | 17.37 | 5.0 | 17.61 |
| SUB_010 | 2.3 | 18.34 | 2.8 | 15.10 | 3.6 | 23.93 | 4.5 | 13.48 | 4.7 | 13.87 |
| SUB_011 | 2.3 | 12.61 | 3.0 | 7.80 | 3.5 | 10.30 | NaN | NaN | NaN | NaN |
| SUB_012 | 2.3 | 35.50 | 2.9 | 25.19 | 3.5 | 38.60 | 4.4 | 18.89 | 4.8 | 20.67 |
| SUB_013 | 2.3 | 40.40 | 3.0 | 29.55 | 3.7 | 43.51 | 4.8 | 19.29 | 5.0 | 19.59 |
| SUB_014 | 2.4 | 21.32 | 3.0 | 13.43 | 3.7 | 22.84 | 4.8 | 10.81 | 5.0 | 10.89 |
| SUB_015 | 2.1 | 38.93 | 2.8 | 22.35 | 3.3 | 30.79 | 4.3 | 8.06 | 4.5 | 8.20 |
| SUB_016 | 2.3 | 35.77 | 3.0 | 25.96 | 3.4 | 27.16 | NaN | NaN | NaN | NaN |
| **Median** | **2.3** | **29.50** | **2.9** | **18.29** | **3.5** | **26.12** | **4.5** | **13.48** | **4.8** | **17.61** |

**Table S3.2** Peak/trough latency (ms) and amplitude (µV) for iTEP by participant at **$SICI_{P1}$ condition**

| SUBJECT | Peak 1 | | Trough 1 | | Peak 2 | | Trough 2 | | Peak 3 | |
|---|---|---|---|---|---|---|---|---|---|---|
| **SUBJECT** | **Lat (ms)** | **Amp (µV)** | **Lat (ms)** | **Amp (µV)** | **Lat (ms)** | **Amp (µV)** | **Lat (ms)** | **Amp (µV)** | **Lat (ms)** | **Amp (µV)** |
| SUB_001 | 2.3 | 26.27 | 3.2 | 2.63 | 3.5 | 3.79 | NaN | NaN | NaN | NaN |
| SUB_002 | 2.4 | 33.88 | 3.1 | 16.58 | 3.7 | 23.67 | NaN | NaN | NaN | NaN |
| SUB_003 | 2.4 | 36.43 | 3.3 | 22.03 | 3.7 | 24.94 | NaN | NaN | NaN | NaN |
| SUB_004 | 2.4 | 18.52 | NaN | NaN | NaN | NaN | NaN | NaN | NaN | NaN |
| SUB_005 | 2.3 | 22.11 | 3.2 | 7.67 | 3.6 | 9.57 | NaN | NaN | NaN | NaN |
| SUB_006 | 2.3 | 37.31 | 3.1 | 14.84 | 3.6 | 18.80 | NaN | NaN | NaN | NaN |
| SUB_007 | 2.5 | 31.89 | 3.4 | 10.82 | 3.9 | 13.27 | NaN | NaN | NaN | NaN |
| SUB_008 | 2.2 | 19.54 | 3.1 | 3.30 | 3.8 | 7.99 | NaN | NaN | NaN | NaN |
| SUB_009 | 2.3 | 22.24 | 3.0 | 16.87 | 3.7 | 27.22 | 4.7 | 17.42 | 5.0 | 18.39 |
| SUB_010 | 2.3 | 18.28 | 2.8 | 14.18 | 3.6 | 23.16 | 4.5 | 13.36 | 4.6 | 13.44 |
| SUB_011 | 2.1 | 8.94 | 3.1 | 0.88 | 3.5 | 1.61 | NaN | NaN | NaN | NaN |
| SUB_012 | 2.3 | 39.87 | 3.1 | 19.74 | 3.6 | 24.79 | NaN | NaN | NaN | NaN |
| SUB_013 | 2.4 | 34.45 | 3.2 | 20.28 | 3.8 | 27.39 | NaN | NaN | NaN | NaN |
| SUB_014 | 2.4 | 22.27 | 3.2 | 8.84 | 3.9 | 13.77 | NaN | NaN | NaN | NaN |
| SUB_015 | 2.2 | 39.21 | 3.1 | 8.93 | 3.5 | 10.89 | NaN | NaN | NaN | NaN |
| SUB_016 | 2.3 | 38.03 | 3.3 | 16.38 | 3.4 | 16.47 | NaN | NaN | NaN | NaN |
| **Median** | **2.3** | **29.08** | **3.1** | **14.18** | **3.6** | **16.47** | **4.6** | **15.39** | **4.8** | **15.91** |

**Table S3.3** Peak/trough latency (ms) and amplitude (μV) for iTEP by participant at **$SICI_{T1}$ condition**

| SUBJECT | Peak 1 | | Trough 1 | | Peak 2 | | Trough 2 | | Peak 3 | |
|---|---|---|---|---|---|---|---|---|---|---|
| **SUBJECT** | **Lat (ms)** | **Amp (μV)** | **Lat (ms)** | **Amp (μV)** | **Lat (ms)** | **Amp (μV)** | **Lat (ms)** | **Amp (μV)** | **Lat (ms)** | **Amp (μV)** |
| SUB_001 | 2.2 | 28.40 | 3.1 | 6.52 | 3.4 | 8.08 | NaN | NaN | NaN | NaN |
| SUB_002 | 2.3 | 33.03 | 3.0 | 18.05 | 3.6 | 24.71 | 4.9 | 4.41 | 5.0 | 4.63 |
| SUB_003 | 2.3 | 34.37 | 3.2 | 23.16 | 3.5 | 24.27 | NaN | NaN | NaN | NaN |
| SUB_004 | 2.3 | 20.11 | NaN | NaN | NaN | NaN | NaN | NaN | NaN | NaN |
| SUB_005 | 2.3 | 22.80 | 3.2 | 9.09 | 3.6 | 11.59 | NaN | NaN | NaN | NaN |
| SUB_006 | 2.2 | 40.91 | 3.1 | 18.97 | 3.5 | 23.23 | NaN | NaN | NaN | NaN |
| SUB_007 | 2.4 | 32.33 | 3.3 | 13.71 | 3.7 | 16.72 | NaN | NaN | NaN | NaN |
| SUB_008 | 2.2 | 16.60 | 3.0 | 5.32 | 3.7 | 9.78 | NaN | NaN | NaN | NaN |
| SUB_009 | 2.3 | 21.65 | 2.9 | 16.09 | 3.6 | 25.82 | 4.6 | 16.82 | 5.1 | 18.70 |
| SUB_010 | 2.3 | 17.12 | 2.8 | 13.85 | 3.6 | 22.83 | 4.5 | 12.89 | 4.6 | 12.96 |
| SUB_011 | 2.3 | 9.87 | 3.1 | 0.91 | 3.8 | 2.05 | NaN | NaN | NaN | NaN |
| SUB_012 | 2.2 | 32.96 | 3.0 | 16.72 | 3.4 | 20.01 | NaN | NaN | NaN | NaN |
| SUB_013 | 2.3 | 42.81 | 3.1 | 27.31 | 3.8 | 37.41 | NaN | NaN | NaN | NaN |
| SUB_014 | 2.4 | 22.43 | 3.1 | 11.08 | 3.7 | 17.62 | NaN | NaN | NaN | NaN |
| SUB_015 | 2.1 | 41.50 | 2.9 | 17.66 | 3.3 | 19.89 | NaN | NaN | NaN | NaN |
| SUB_016 | 2.3 | 42.39 | NaN | NaN | NaN | NaN | NaN | NaN | NaN | NaN |
| **Median** | **2.3** | **30.37** | **3.1** | **14.97** | **3.6** | **19.95** | **4.6** | **12.89** | **5.0** | **12.96** |

**Table S3.4** Peak/trough latency (ms) and amplitude (µV) for iTEP by participant at **$SICI_{P2}$ condition**

| SUBJECT | Peak 1 | | Trough 1 | | Peak 2 | | Trough 2 | | Peak 3 | |
|---|---|---|---|---|---|---|---|---|---|---|
| SUBJECT | Lat (ms) | Amp (µV) | Lat (ms) | Amp (µV) | Lat (ms) | Amp (µV) | Lat (ms) | Amp (µV) | Lat (ms) | Amp (µV) |
| SUB_001 | 2.2 | 31.86 | 3.1 | 9.83 | 3.4 | 11.25 | NaN | NaN | NaN | NaN |
| SUB_002 | 2.3 | 33.87 | 3.0 | 20.56 | 3.5 | 26.15 | 4.6 | 7.40 | 4.7 | 7.49 |
| SUB_003 | 2.3 | 33.20 | 3.2 | 20.77 | 3.4 | 21.09 | NaN | NaN | NaN | NaN |
| SUB_004 | 2.3 | 19.72 | NaN | NaN | NaN | NaN | NaN | NaN | NaN | NaN |
| SUB_005 | 2.3 | 23.14 | 3.0 | 10.17 | 3.5 | 13.43 | NaN | NaN | NaN | NaN |
| SUB_006 | 2.2 | 41.42 | 3.0 | 21.18 | 3.5 | 25.98 | NaN | NaN | NaN | NaN |
| SUB_007 | 2.3 | 31.38 | 3.2 | 14.27 | 3.8 | 18.10 | NaN | NaN | NaN | NaN |
| SUB_008 | 2.1 | 22.16 | 3.0 | 9.16 | 3.5 | 13.36 | NaN | NaN | NaN | NaN |
| SUB_009 | 2.3 | 22.16 | 2.9 | 17.70 | 3.7 | 27.99 | 4.5 | 19.55 | 4.9 | 21.43 |
| SUB_010 | 2.3 | 20.18 | 2.8 | 15.74 | 3.5 | 24.79 | 4.4 | 14.24 | 4.7 | 14.68 |
| SUB_011 | 2.3 | 9.40 | 3.1 | 2.06 | 3.5 | 3.24 | NaN | NaN | NaN | NaN |
| SUB_012 | 2.3 | 36.06 | 3.1 | 13.98 | 3.7 | 18.67 | NaN | NaN | NaN | NaN |
| SUB_013 | 2.3 | 37.72 | 3.1 | 22.61 | 3.7 | 30.94 | NaN | NaN | NaN | NaN |
| SUB_014 | 2.3 | 23.37 | 3.1 | 11.03 | 3.7 | 16.73 | NaN | NaN | NaN | NaN |
| SUB_015 | 2.1 | 40.79 | 2.9 | 17.29 | 3.3 | 20.72 | NaN | NaN | NaN | NaN |
| SUB_016 | 2.3 | 37.02 | NaN | NaN | NaN | NaN | NaN | NaN | NaN | NaN |
| **Median** | **2.3** | **31.62** | **3.1** | **15.01** | **3.5** | **19.69** | **4.5** | **14.24** | **4.7** | **14.68** |

**Effect of paired pulse on iTEP peak 3 reflected in GMFA and local iTEP waveforms**

Figure S5A shows the relationship between inhibition of iTEP Peak3, expressed as GMFA-AUC ratio, and corticomotor inhibition, expressed the MEP ratio in the **FDI** hand muscle. SUB001 (marked by dashed red line) showed marked paired-pulse GMFA facilitation. Visual inspection suggests, within the Peak3 time window, GMFA value was greater in the paired-pulse conditions than in the single-pulse TS condition, whereas iTEP amplitudes at a channel near the stimulation site (channel 16) were reduced (Figure S5B). This suggests a divergence between paired-pulse effects on global response strength, indexed by GMFA, and on local iTEP morphology at an electrode near the stimulation site for the peak3 time window.

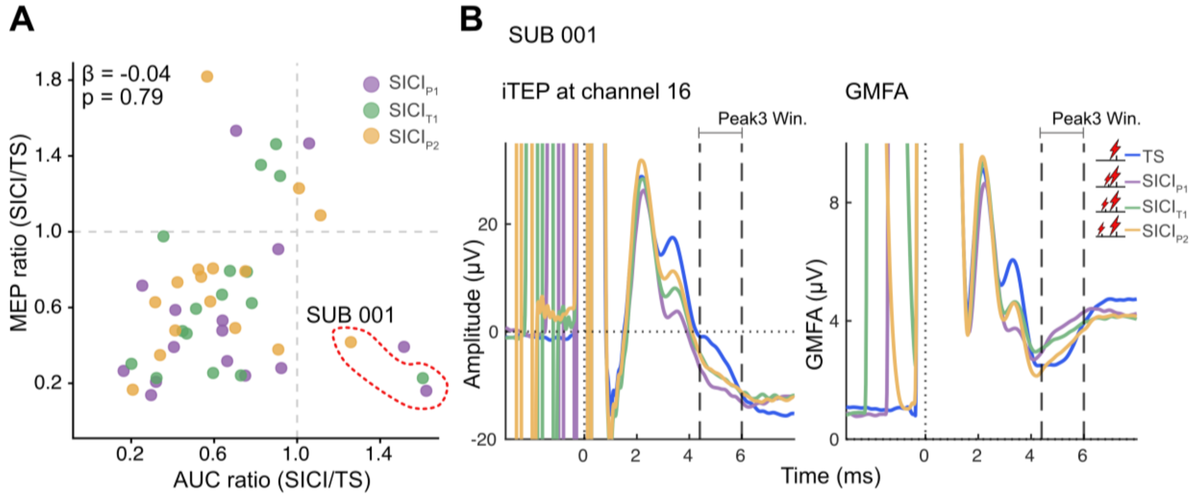


**Figure S5.** Relationship between cortical and corticomotor inhibition in the main experiment. **A.** Paired-pulse (SICI$_{P1}$, SICI$_{T1}$, and SICI$_{P2}$) to single-pulse TS AUC ratio in the iTEP Peak3 time window, plotted against the corresponding paired-pulse to single-pulse MEP ratio in hand muscle **FDI**. ISIs in SICI$_{P1}$, SICI$_{T1}$, and SICI$_{P2}$ were adjusted to Peak 1 (P1), Trough 1 (T1), and Peak 2 (P2) of the individual SICF curve, respectively. The regression coefficient β and corresponding p-value are noted in the upper-left corner. **B.** iTEPs (left) at a channel close to the site of stimulation (channel 16) and GMFA (right) from SUB001 (demarcated by dashed red line in A) for the single-pulse condition (TS) and the three paired-pulse SICI conditions (SICI$_{P1}$, SICI$_{T1}$, SICI$_{P2}$). ISIs in SICI$_{P1}$, SICI$_{T1}$, and SICI$_{P2}$ were adjusted to Peak 1 (P1), Trough 1 (T1), and Peak 2 (P2) of the individual SICF curve, respectively.

AUC = area under curve, GMFA= Global Mean Field Amplitude, ISI = interstimulus interval, SICI = short-latency intracortical inhibition, SICF= short-latency intracortical facilitation, TS = Test Stimulus.

**Effects of paired pulse on ADM MEP amplitude in the main experiment**

Compared with TS, the three paired-pulse SICI conditions ($SICI_{P1}$, $SICI_{T1}$, $SICI_{P2}$) resulted in smaller MEP amplitudes in the hand muscle ADM (Figure S6). This was supported by a significant main effect of condition on log-transformed MEP amplitude ($F(3, 42) = 12.34$, $p < 0.0001$). Post-hoc comparisons showed that each SICI condition produced smaller MEPs than the single-pulse TS ($SICI_{P1}$ vs. TS: p = <0.0001; $SICI_{T1}$ vs. TS: p = 0.001; $SICI_{P2}$ vs. TS: p = 0.023). In addition, the shortest-ISI SICI condition yielded smaller MEPs than the longest-ISI condition ($SICI_{P1}$ vs $SICI_{P2}$: p = 0.015).

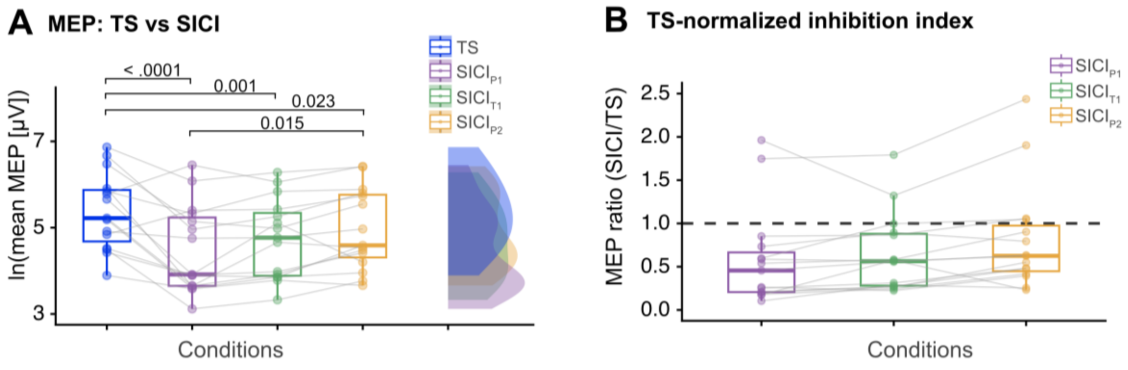


**Figure S6**. Paired-pulse MEP inhibition in the main experiment. A. Mean MEP amplitudes from the single-pulse condition (TS) and the three paired-pulse SICI conditions ($SICI_{P1}$, $SICI_{T1}$, $SICI_{P2}$) amplitude in hand muscle **ADM**. Natural-log-transformed values are shown as boxplots and density plots, with overlaid points for individual subjects (**n=15**) and grey lines indicating within-subject changes. ISIs in $SICI_{P1}$, $SICI_{T1}$, and $SICI_{P2}$ were adjusted to Peak 1 (P1), Trough 1 (T1), and Peak 2 (P2) of the individual SICF curve, respectively. Significant differences in post-hoc tests are indicated with horizontal lines with Holm-adjusted p-values noted above. B. Relative MEP inhibition, expressed as the ratio of paired-pulse ($SICI_{P1}$, $SICI_{T1}$, and $SICI_{P2}$) to single-pulse TS MEP amplitude, shown as boxplots with overlaid points for individual subjects and grey lines indicating within-subject changes. The dashed horizontal line at 1 marks the TS reference level.

ISI = interstimulus interval, SICI = short-latency intracortical inhibition, SICF= short-latency intracortical facilitation, TS = Test Stimulus.

**Cortical inhibition does not covary with inhibition of MEP in hand muscle ADM in the main experiment**

Cortical inhibition was indexed by AUC_ratio for iTEP Peak2, Peak3, and N15. To examine their relationships with corticomotor inhibition indexed by MEP ratio in hand muscle ADM, we fitted separate linear mixed-effects models relating AUC_ratio to MEP_ratio, separately for Peak2, Peak3, and N15. Likelihood-ratio tests provided no evidence that the AUC-MEP association differed across SICI ISIs. We therefore report the additive models. None of the AUC ratios showed a significant association with the MEP ratio (Figure S7).

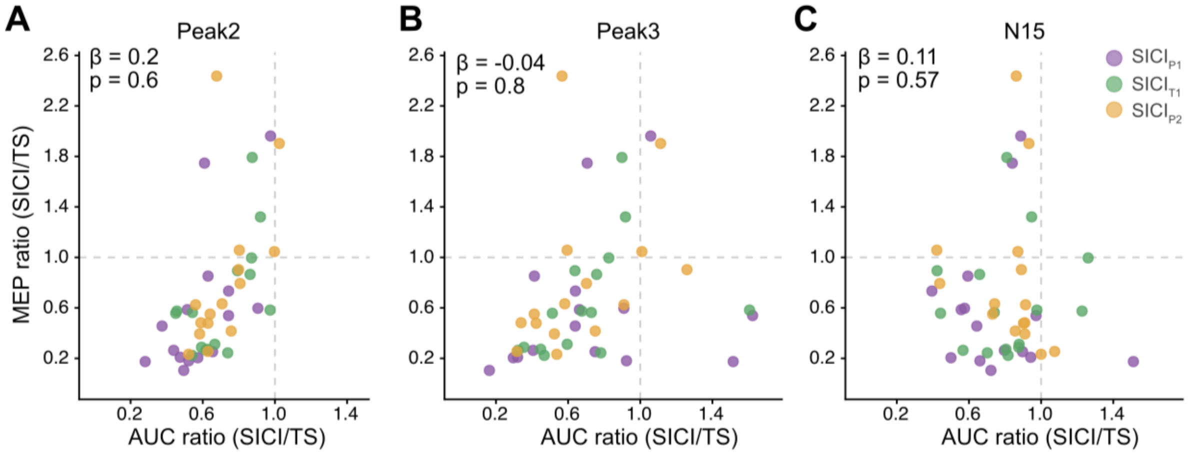


**Figure S7**. Relationship between cortical and corticomotor inhibition in the main experiment for ADM. A. Paired-pulse ($SICI_{P1}$, $SICI_{T1}$, and $SICI_{P2}$) to single-pulse TS AUC ratio in the iTEP Peak2 time window, plotted against the corresponding paired-pulse to single-pulse MEP ratio in hand muscle **ADM**. ISIs in $SICI_{P1}$, $SICI_{T1}$, and $SICI_{P2}$ were adjusted to Peak 1 (P1), Trough 1 (T1), and Peak 2 (P2) of the individual SICF curve, respectively. The regression coefficient β and corresponding p-value are noted in the upper-left corner. B. Same as in A, except that the paired-pulse to single-pulse TS AUC ratio was computed for iTEP Peak3 time window. C. Same as in A, except that the paired-pulse to single-pulse TS AUC ratio was computed for the TEP N15 time window.

AUC = area under curve, ISI = interstimulus interval, SICI = short-latency intracortical inhibition,
